# Reduced progranulin increases tau and *α*-synuclein inclusions and alters phenotypes of tauopathy mice via glucocerebrosidase

**DOI:** 10.1101/2022.12.25.521308

**Authors:** Hideyuki Takahashi, Sanaea Bhagwagar, Sarah H. Nies, Marius T. Chiasseu, Guilin Wang, Ian R. Mackenzie, Stephen M. Strittmatter

## Abstract

Comorbid proteinopathies are observed in many neurodegenerative disorders including Alzheimer’s disease (AD), increase with age, and influence clinical outcomes, yet the mechanisms remain ill-defined. Here, we show that reduction of progranulin (PGRN), a lysosomal protein associated with TDP-43 proteinopathy, also increases tau inclusions, causes concomitant accumulation of *α*-synuclein and worsens mortality and disinhibited behaviors in tauopathy mice. The increased inclusions paradoxically protect against spatial memory deficit and hippocampal neurodegeneration. PGRN reduction with tau pathology attenuates activity of *β*-glucocerebrosidase (GCase), a protein previously associated with synucleinopathy, while increasing glucosylceramide (GlcCer)-positive tau inclusions. In neuronal culture, GCase inhibition enhances tau aggregation induced by AD-tau. Furthermore, purified GlcCer directly promotes tau aggregation *in vitro*. Neurofibrillary tangles in human tauopathies are also GlcCer-immunoreactive. Thus, in addition to TDP-43, PGRN regulates tau- and synucleinopathies via GCase and GlcCer. A lysosomal PGRN–GCase pathway may be a common therapeutic target for age-related comorbid proteinopathies.

## Introduction

Many neurodegenerative disorders are characterized by abnormal accumulation of protein aggregates in the brain. Alzheimer’s disease (AD) and frontotemporal lobar degeneration (FTLD) are characterized by tau and/or TAR DNA-binding protein 43 (TDP-43) inclusions while *α*-synuclein (*α*-syn) accumulates in the brain of Parkinson’s disease (PD) and dementia with Lewy bodies (DLB) patients ^1^. In addition to these primary proteinopathies defining the diseases, it is also known that additional proteinopathies often accumulate as co-pathologies in many neurodegenerative diseases. For example, accumulation of *α*-syn is frequently seen in AD cases. These concomitant proteinopathies increase with age and affect clinical course, therefore have implications for clinical trials targeting only a single disease-associated protein. Global decline in machinery to maintain cellular proteostasis, cross-seeding, and genetic factors are hypothesized to account for the comorbidities, but the exact molecular mechanisms are not fully understood ^2–4^.

Progranulin (PGRN), encoded by the *GRN* gene in humans, is a secreted and lysosomal glycoprotein that plays an important role in lysosomal homeostasis. PGRN is mainly produced by neurons and microglia in the CNS and is involved in several neurodegenerative disorders associated with lysosomal dysfunction ^5–7^. While heterozygous *GRN* mutations are a frequent cause of familial FTLD ^8, 9^, rare homozygous *GRN* mutations cause the lysosomal storage disorder neuronal ceroid lipofuscinosis ^10^. Several *GRN* polymorphisms are associated with increased risk for Gaucher disease, a common lysosomal storage disorder ^11^.

Although PGRN was initially linked to FTLD with TDP-43 inclusions, *GRN* mutations are also reported in a substantial number of AD and PD patients ^12–19^. In addition, genetic studies have suggested that *GRN* variations increases a risk for AD and PD ^20–25^. A *GRN* AD risk variant is associated with increased cerebrospinal fluid tau levels^26^. Accumulation of tau and/or *α*-syn in addition to TDP-43 inclusions is observed in several FTLD patients with different *GRN* mutations ^16, 27–32^. In preclinical models, we and others have found an increase in tau pathology in *Grn*^−/−^ mice injected with AAV-human P301L tau and P301L tau transgenic mice with PGRN reduction ^26, 33^. These studies suggest that lysosomal PGRN regulates not only TDP-43 but also other proteinopathies, especially tau- and synucleinopathy, and therefore may be a common therapeutic target for multiple proteinopathies in neurodegenerative diseases. However, the mechanisms by which PGRN regulates other proteinopathies and whether PGRN reduction affects their symptoms are currently not clear.

In this study, to gain insights into the relationship between PGRN and tauopathy, we analyze the well-characterized PS19 tauopathy mouse model overexpressing human P301S tau ^34^ on PGRN haploinsufficient and complete null backgrounds ^35^. Here, we show that both complete loss and haploinsufficiency of PGRN increase tau inclusions, cause concomitant accumulation of *α*-syn, and worsen mortality and disinhibited behaviors in PS19 mice. Reduction of PGRN paradoxically protects against a spatial memory deficit and hippocampal neurodegeneration and transcriptomic change in PS19 mice. We find that PGRN reduction with tauopathy significantly impairs activity of *β*-glucocerebrosidase (GCase), a protein previously associated with synucleinopathy ^36, 37^, while increasing tau inclusions that are immunoreactive for GCase substrate glucosylceramide (GlcCer). In neuronal culture, GCase inhibition promotes tau aggregation induced by AD brain-derived tau fibrils. *In vitro* studies show that purified GlcCer directly accelerates tau aggregation. Neurofibrillary tangles in human tauopathy brains are also immunoreactive for GlcCer. Thus, our study reveals unexpected role of GlcCer in tau aggregation and demonstrates that PGRN regulates formation of tau and *α*-syn inclusion via GCase to alter symptoms and neurodegeneration. A lysosomal PGRN–GCase pathway may have therapeutic potential in comorbid proteinopathies.

## Results

### PGRN reduction decreases body weight and survival rate and worsens disinhibited behavior of PS19 mice

To assess effects of PGRN reduction on tauopathy, we crossed PS19 and *Grn*^−/−^ mice and generated 6 genotypes (WT, *Grn*^+/−^, *Grn*^−/−^, PS19, PS19 *Grn*^+/−^, and PS19 *Grn*^−/−^ mice). We first monitored their body weights and found a significant decrease in body weight of PS19 *Grn*^−/−^ mice, but not the other genotypes, compared to WT mice at 7.5 months of age (Fig. 1a). A significant decrease in the body weight was observed in PS19 mice at 9 months of age. Death due to hindlimb paresis observed in PS19 mice ^34, 38^ was also slightly increased in PS19 *Grn*^−/−^ mice at 10 months of age (Fig. 1b).

**Fig. 1:**
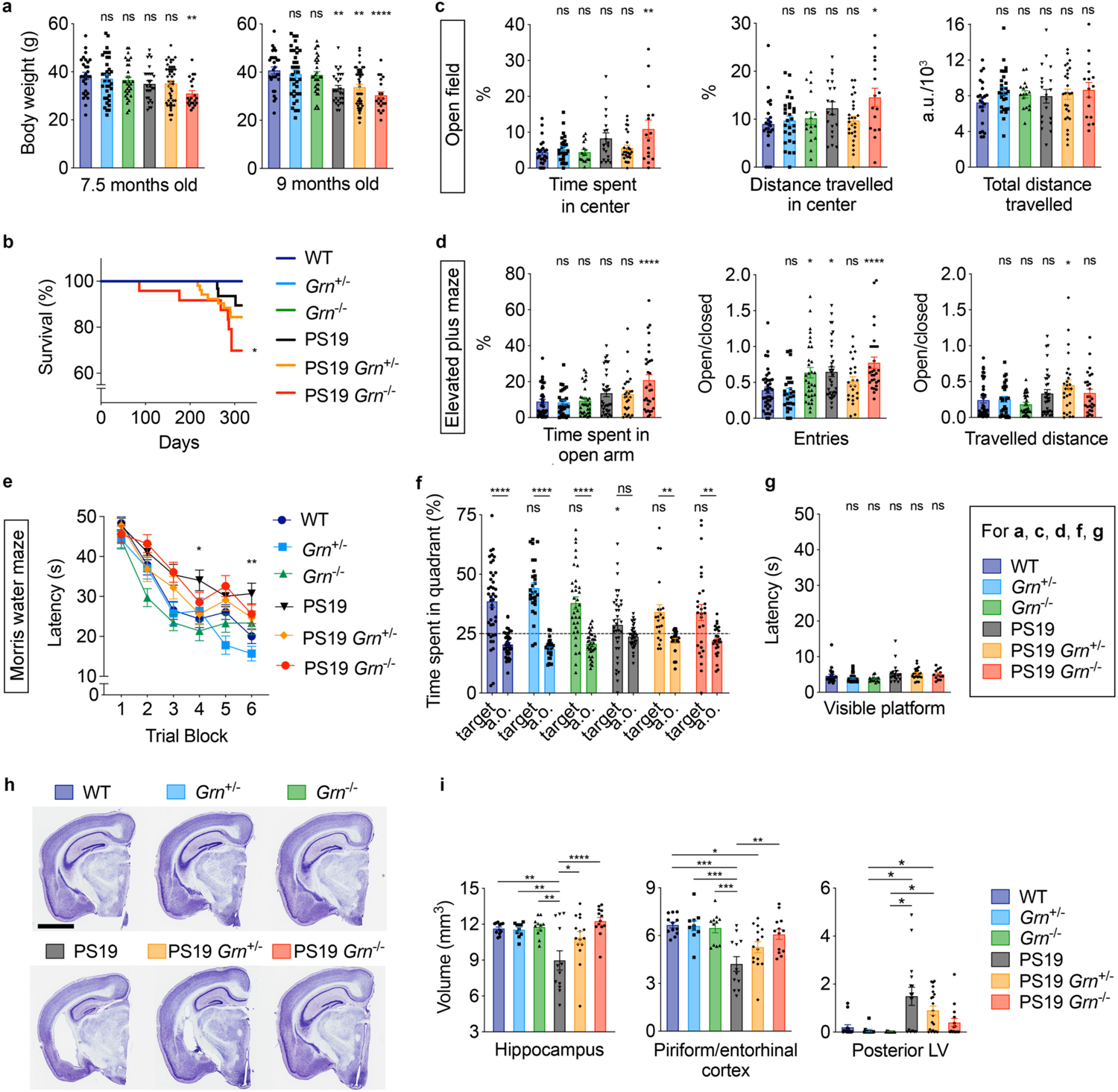
PGRN reduction worsens mortality and disinhibition while improving a memory deficit and hippocampal atrophy in PS19 mice. **a.** Body weight of 6 genotypes (WT, *Grn*^+/−^, *Grn*^−/−^, PS19, PS19 *Grn*^+/−^, and PS19 *Grn*^−/−^ mice) at 7.5 and 9 months of age. Mean ± SEM, n = 20–47 mice per genotype, **p < 0.01; ****p < 0.0001; One-way ANOVA with Dunnett’s post hoc test. **b.** Survival curves of 6 genotypes. WT, n = 36; *Grn*^+/−^, n = 57; *Grn*^−/−^, n = 30; PS19, n = 31; PS19 *Grn*^+/−^, n = 53; PS19 *Grn*^−/−^, n = 24. *p < 0.05; Versus WT, pairwise log-rank test. **c.** Open field test of 6 genotypes at 10-11 months of age showing time spent in the center zone (%), distance travelled in the center zone (%), and total distance travelled during the test. Mean ± SEM, n = 16–28 mice per genotype, *p < 0.05; **p < 0.01; One-way ANOVA with Dunnett’s post hoc test. **d.** Elevated plus maze (EPM) of 6 genotypes at 10-11 months of age showing time spent in the open arms (%), ratio of number of entries into the open versus closed arms, and ratio of distance travelled in the open versus closed arms of the maze. Mean ± SEM, n = 27–41 mice per genotype, *p < 0.05; ****p < 0.0001; One-way ANOVA with Dunnett’s post hoc test. **e.** Morris water maze (MWM) learning trial of 6 genotypes at 10-11 months of age. Spatial learning is plotted as latency to find hidden platform. Mean ± SEM, WT, n = 21–41 mice per genotype; In the trial block 4 and 6, the PS19 group differed from the WT group, whereas none of the other groups differed from the WT group, as shown in figure. *p < 0.05, **p < 0.01; One-way ANOVA, with Dunnett’s post hoc test. **f.** MWM probe trial of 6 genotypes at 10-11 months of age. Percentage of time spent in the target quadrant and averaged time spent in all other (a.o.) quadrants. Mean ± SEM, n = 21–41 mice per genotype. *p < 0.05, **p < 0.01, ****p < 0.0001; Top: T test with Welch’s correction. Bottom: One-way ANOVA, with Dunnett’s post hoc test comparing to WT (only using target quadrants). Additionally, one sample t test showed that the mean time of PS19 mice, but not the others, in the target quadrant is not different from random chance performance of 25% (dashed line) (p = 0.139). **g.** Visible platform trial of 6 genotypes at 10-11 months of age. n = 13–28 mice per genotype. **h.** Representative images of Nissl staining of sections from 6 genotypes (WT, *Grn*^+/−^, *Grn*^−/−^, PS19, PS19 *Grn*^+/−^, and PS19 *Grn*^−/−^ mice). Bar, 2 mm. **i.** Volumes of the hippocampus, piriform/entorhinal cortex, and posterior lateral ventricle (LV) in 6 genotypes at 9-12 months of age. Mean ± SEM, n = 10–15 mice (the hippocampus and piriform/entorhinal cortex) and 12–20 mice (the posterior LV) per genotype, *p < 0.05; **p < 0.01; ***p < 0.001; ****p < 0.0001; One-way ANOVA with Tukey’s post hoc test or Kruskal-Wallist test with Dunn’s post hoc test (for the posterior LV).

Open field tests with the remaining cohort at 10-11 months of age showed significant increases in permanence time and travelled distance in center area of the field for PS19 *Grn*^−/−^ mice, but not the other genotypes, compared to WT mice, while total distance travelled was similar between the 6 genotypes, suggesting an exacerbation of disinhibition in PS19 *Grn*^−/−^ mice (Fig. 1c). In line with the results of open field test, in elevated plus maze (EPM), PS19 *Grn*^−/−^ mice, but not the other genotypes, spent more time in the open arms, compared to WT mice. Consistent with previous studies ^39, 40^, PS19 and *Grn*^−/−^ mice also entered to the open arms more frequently than WT mice, but the frequency was further exacerbated in PS19 *Grn*^−/−^ mice. In addition, a greater travelled distance in the open arms was observed for PS19 *Grn*^+/−^ mice (Fig. 1d). Similar results were also observed when only male animals were analyzed (Extended Data Fig. 1a). These results suggest that PGRN reduction exacerbates disinhibited behavior in PS19 mice.

### PGRN reduction improves a memory impairment and hippocampal neurodegeneration in PS19 mice

To assess spatial learning and memory of the 6 genotypes, we also tested the same cohorts in Morris water maze (MWM). Consistent with previous studies ^40, 41^, PS19 mice showed significant learning and memory impairments in the MWM paradigm (Fig. 1e,f). Unexpectedly, in learning trials, the latency to find the platform in PS19 *Grn*^+/−^ or PS19 *Grn*^−/−^ mice was indistinguishable from that of WT mice in trial blocks 4 and 6. In a probe trial after the learning trials, PS19 *Grn*^+/−^ or PS19 *Grn*^−/−^ mice spent significantly more time in the target quadrant than average of all other quadrants, suggesting improved memory retention in these mice (Fig. 1f). The latency was similar between the 6 genotypes in the visible platform trial (Fig. 1g). Similar results in MWM were also observed when only male animals were analyzed (Extended Data Fig. 1b,c). Thus, PGRN reduction improved a memory impairment while exacerbating disinhibited behavior in PS19 mice.

To provide insight into the basis for the behavioral results, we assessed neurodegeneration and pathological changes in the brains of the 6 genotypes. Previous studies reported a significant difference between male and female in neurodegeneration and tau pathology in PS19 mice ^42–45^. Thus, subsequent analyses were performed using only male animals at 9-12 months of age unless otherwise noted. We first analyzed PGRN levels in PS19 mice using immunohistochemistry and found an increase in PGRN immunoreactivity, which was primarily associated with microgliosis, in the brain of PS19 mice. A reduction and loss of PGRN immunoreactivity were confirmed in PS19 *Grn*^+/−^ and PS19 *Grn*^−/−^ mice, respectively (Extended Data Fig. 2).

**Fig. 2:**
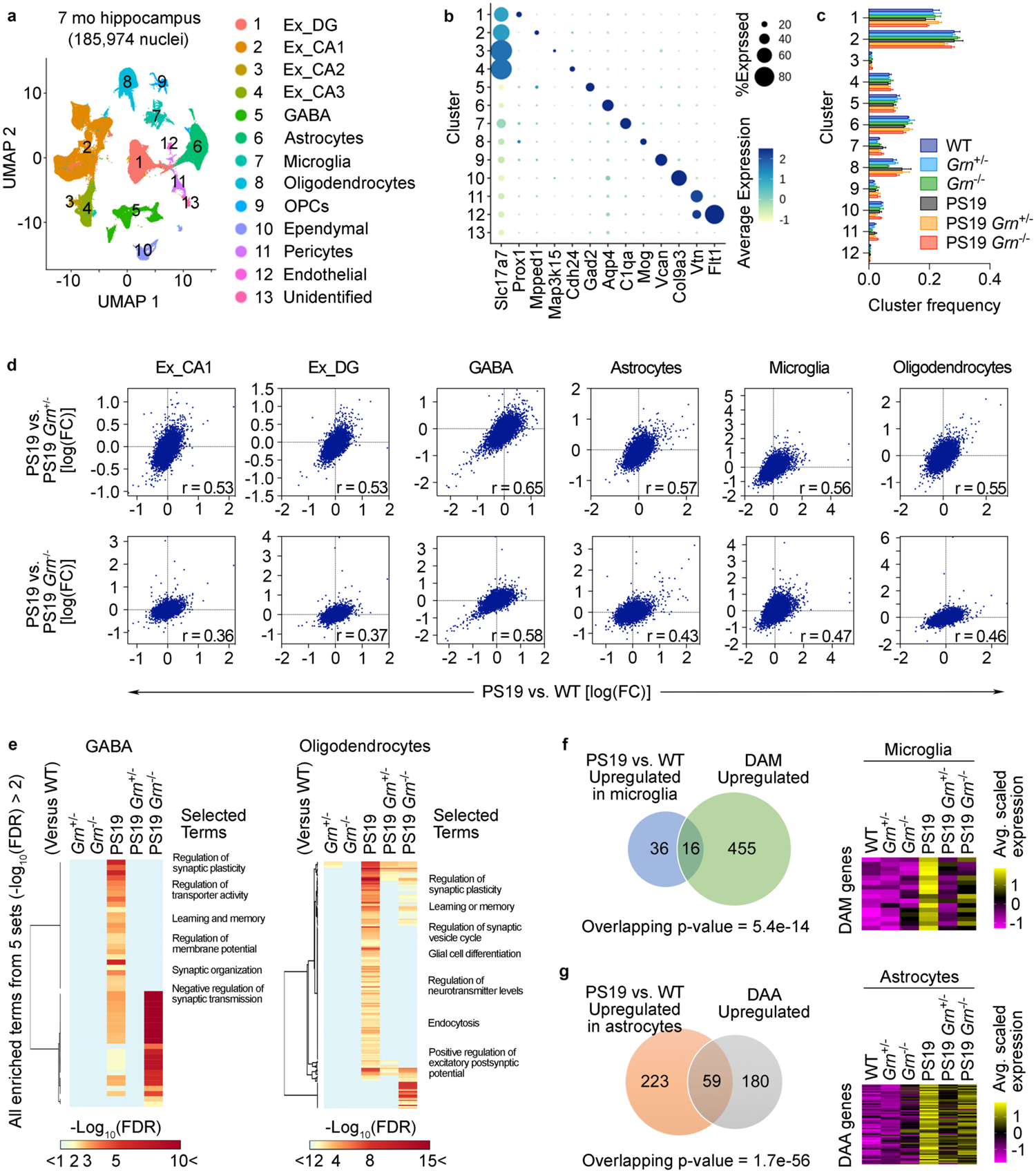
PGRN reduction attenuates transcriptomic changes in PS19 mice. **a.** UMAP plot showing 13 cell type clusters of 185,974 single-nucleus RNA profiles from hippocampi of 6 genotypes (WT, *Grn*^+/−^, *Grn*^−/−^, PS19, PS19 *Grn*^+/−^, and PS19 *Grn*^−/−^ mice) with three biological replicates at 7 months of age. **b.** Dot plot showing the percent of cells expressing marker genes and average expression across all cell type clusters. **c.** Quantification of frequency of each cluster in 6 genotypes. No significant difference was observed between 6 genotypes in each cluster (One-way ANOVA). **d.** Transcriptome-wide rescue/exacerbation plots showing the relationship between log(FC)s of genes in PS19 vs. WT and PS19 vs. PS19 *Grn*^+/−^ or PS19 *Grn*^−/−^ mice in the indicated cell clusters. Genes that express > 1% of all nuclei in either genotype were used. The correlation coefficients (r) are indicated in the plots. All correlations were significant (p > 1e-15). **e.** Heatmaps showing values of –log10(FDR) of all enriched terms identified from 5 comparisons (*Grn*^+/−^, *Grn*^−/−^, PS19, PS19 *Grn*^+/−^, or PS19 *Grn*^−/−^ vs. WT) (defined by – log_10_(FDR) > 2) in the same 5 comparisons in GABA and Oligodendrocytes cell type clusters. Selected terms are highlighted. **f.** Effects of PGRN reduction on tau-induced expression of DAM-associated genes. Venn diagram shows the overlap between DEGs in PS19 microglia (vs. WT) and DAM-associated genes. Heatmap shows expression of the overlapping genes in microglia of 6 genotypes. Overlapping p-values was calculated by Fisher’s exact test. **g.** Effects of PGRN reduction on tau-induced expression of DAA-associated genes. Venn diagram shows the overlap between DEGs in PS19 astrocytes (vs. WT) and DAA-associated genes. Heatmap shows expression of the overlapping genes in astrocytes of 6 genotypes. Overlapping p-values was calculated by Fisher’s exact test.

We then investigated whether PGRN reduction affects brain atrophy, enlarged posterior lateral ventricle, and hippocampal neurodegeneration in PS19 mice ^34, 38, 43, 44, 46^. Strikingly, PGRN reduction significantly attenuated atrophy of the hippocampus and piriform/entorhinal cortex of PS19 mice in a gene dosage-dependent manner (Fig. 1h,i). A significant increase in the posterior lateral ventricle was observed in PS19 mice compared to *Grn*^+/−^ or *Grn*^−/−^ mice but the increase was absent in PS19 *Grn*^−/−^ mice (Fig. 1h,i). Hippocampal CA1 pyramidal neuronal and dentate gyrus (DG) granule cell layers were thinner in PS19, but not in PS19 *Grn*^+/−^, or PS19 *Grn*^−/−^ mice compared to WT mice (Extended Data Fig. 1d,e). These results revealed an unexpected protective role for PGRN reduction in tau-mediated brain atrophy and hippocampal neurodegeneration.

### PGRN reduction attenuates hippocampal transcriptomic changes in PS19 mice

To further characterize the protective role for PGRN reduction in neurodegeneration at transcriptomic levels, we performed single-nucleus RNA sequencing (snRNA-seq) with nuclei isolated from hippocampi of the 6 genotypes with three biological replicates (18 mice, 185,974 nuclei). We used 7-month-old male animals for this analysis to gain insights into earlier changes, rather than late secondary effects due to neurodegeneration. RNA expression profiles of all nuclei were visualized by Uniform Manifold Approximation and Projection (UMAP) and grouped into 13 cell type clusters based on expression of known marker genes (Fig. 2a,b, and Extended Data Fig. 3a). Fractions of each cell type cluster were not significantly different between the 6 genotypes (Fig. 2c), suggesting that there is no significant neurodegeneration or gliosis in PS19 mice at this age.

**Fig. 3:**
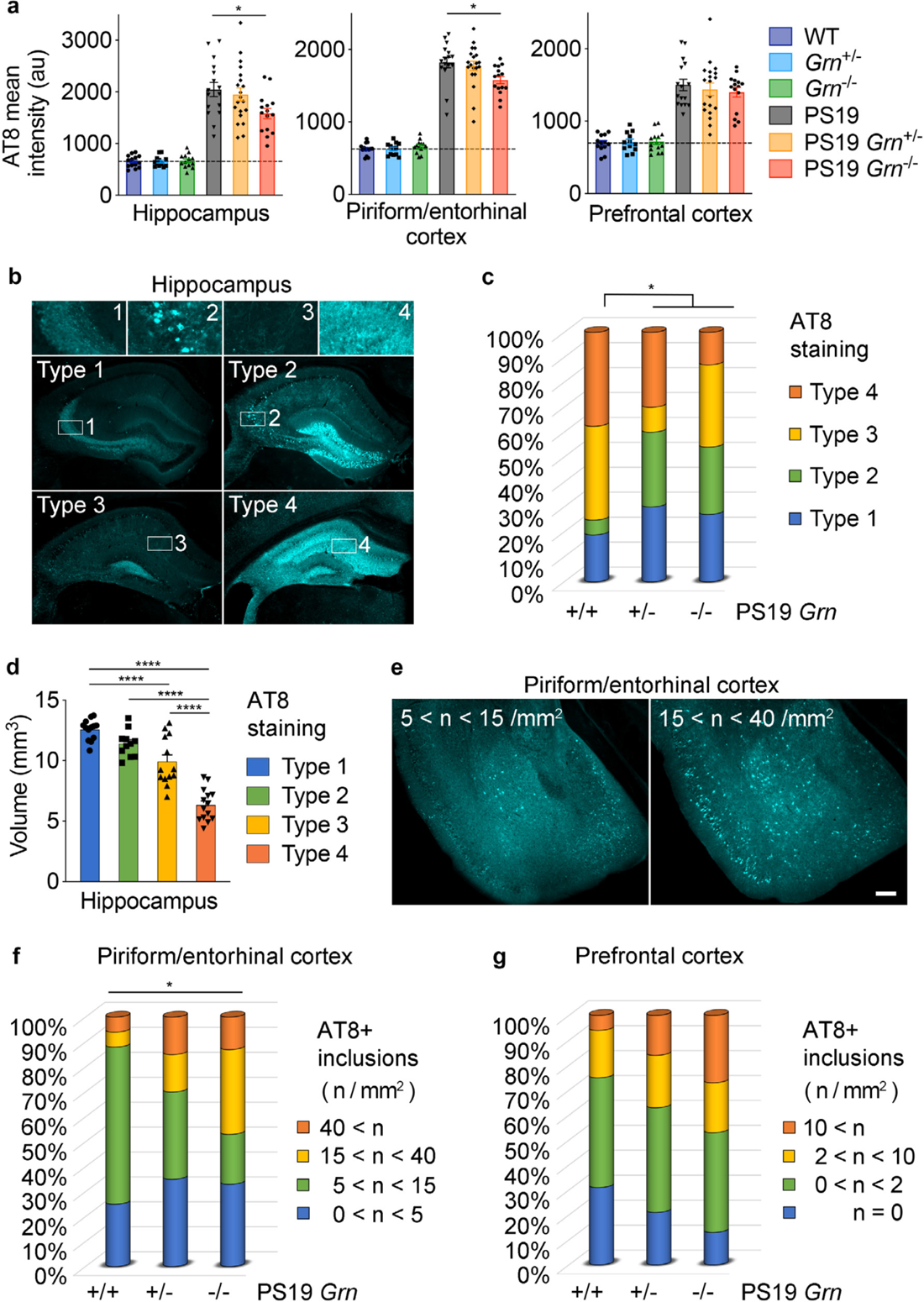
PGRN reduction alters AT8 staining pattern and increases tau inclusions in PS19 mice. **a.** AT8 mean intensity of the hippocampus, piriform/entorhinal cortex, and prefrontal cortex of 6 genotypes (WT, *Grn*^+/−^, *Grn*^−/−^, PS19, PS19 *Grn*^+/−^, and PS19 *Grn*^−/−^ mice) at 9-12 months of age. Mean ± SEM, n = 12–20 mice per genotype, *p < 0.05; One-way ANOVA with Tukey’s post hoc test (in PS19, PS19 *Grn*^+/−^, and PS19 *Grn*^−/−^ mice). **b.** Representative images of hippocampal four distinct AT8 staining types. **c.** Distribution of the AT8 staining types in 6 genotypes at 9-12 months of age. n = 15–20 mice per genotype, *p < 0.05; Fisher’s exact test (2 rows (PS19 versus PS19 *Grn*^+/−^ + PS19 *Grn*^−/−^) and 2 columns (type 1+2 versus type 3+4)) **d.** Association of hippocampal atrophy with four AT8 staining types. Mean ± SEM, n = 51 mice, ****p < 0.0001; One-way ANOVA with Tukey’s post hoc test. **e.** Representative images of AT8 staining in the piriform/entorhinal cortex showing density of 5 < n < 15 inclusions /mm^2^ or 15 < n < 40 inclusions /mm^2^. Bar, 200 μm. **f.** Distribution of the number of AT8-positive inclusions in the piriform/entorhinal of 6 genotypes at 9-12 months of age. n = 15–20 mice per genotype, *p < 0.05; Chi-square test for trend (3 rows (PS19, PS19 *Grn*^+/−^, versus PS19 *Grn*^−/−^) and 2 columns (0 < n < 15 versus 15 < n)) **g.** Distribution of the number of AT8-positive inclusions in the prefrontal cortex of 6 genotypes at 9-12 months of age. n = 15–20 mice per genotype.

Differential expression analysis in each cluster identified multiple differentially-expressed genes (DEGs) in PS19 nuclei compared to WT, particularly in DG granule cells (Ex_DG), CA1 pyramidal (Ex_CA1), GABAergic (GABA), astrocytes, and oligodendrocytes cell type clusters. In addition, we found significant gene overlaps between DEGs from PS19 versus WT and from PS19 versus PS19 with PGRN reduction for several cell type clusters including excitatory neurons, GABAergic neurons, and oligodendrocytes (Extended Data Fig. 3c). Across cell types, substantially fewer DEGs were detected for non-PS19 mice when comparing *Grn*^+/−^ and *Grn*^−/−^ to WT (Extended Data Fig. 3d,e). Overall, the results show that mutant tau causes transcriptomic changes in PS19 neurons and glia, and these are attenuated by PGRN reduction. In fact, a transcriptome-wide rescue effect ^47^ was also observed in most cell types of PS19 mice with PGRN reduction (Fig. 2d). Please note that a slope of 1 in these correlation plots reflects full rescue of DEGs. Furthermore, enrichment analysis focusing on GABAergic and oligodendrocyte clusters revealed that PGRN reduction significantly decreases enriched terms identified from DEGs in PS19 versus WT (Fig. 2e). Recent single cell RNA-seq analyses have identified disease-associated microglia (DAM) and astrocytes (DAA) signatures in mouse models of AD ^48, 49^. Upregulated genes in PS19 microglia and astrocytes partly overlapped with those upregulated in DAM and DAA (Fig. 2f,g). Importantly, PGRN reduction also decreased expression of these genes (Fig. 2f,g). Together, PGRN reduction attenuated neuronal and glial transcriptomic changes in the hippocampus of PS19 mice.

### PGRN reduction alters tau pathology pattern and increases tau inclusions

The rescue of brain atrophy and global gene expression in PS19 *Grn*^+/−^ and PS19 *Grn*^−/−^ hippocampi suggests an effect of PGRN reduction on tau pathology. Therefore, we next examined tau pathology of PS19 mice with PGRN reduction. AT8 (p-S202/T205 tau) immunostaining showed a mild decrease in AT8 mean intensity in the hippocampus and piriform/entorhinal cortex of PS19 *Grn*^−/−^, but not in PS19 *Grn*^+/−^ mice (Fig. 3a). We further analyzed four distinct AT8 staining types in our PS19 *Grn*^+/−^ and *Grn*^−/−^ mice by a previously reported method ^43, 46, 50^. Consistent with the previous studies, four AT8 staining types were correlated with hippocampal atrophy in our PS19 cohorts (Fig. 3b,d). Intriguingly, while PGRN reduction decreases the percentage of type 4 in PS19 mice, we observed a slight increase in type 1, and also a significant increase in type 2, which is characterized by tangle-like cell body staining, in PS19 mice with PGRN reduction (Fig. 3c). In the piriform/entorhinal and prefrontal cortex of PS19 mice with PGRN reduction, we observed an increase in the fraction of mice with a high density of inclusions (Fig. 3e-g). Immunoblot analysis using anti-total tau antibody of the formic-acid-soluble fraction showed that the increase in tau inclusions was due to subcellular distribution and not attributable to overall changes in insoluble tau in PS19 mice with PGRN reduction (Extended Data Fig. 4e). These results suggest that PGRN reduction accelerates formation of tangle-like cell body tau inclusions in PS19 mice, which is partly consistent with previous studies using different tauopathy models^26, 33^.

**Fig. 4:**
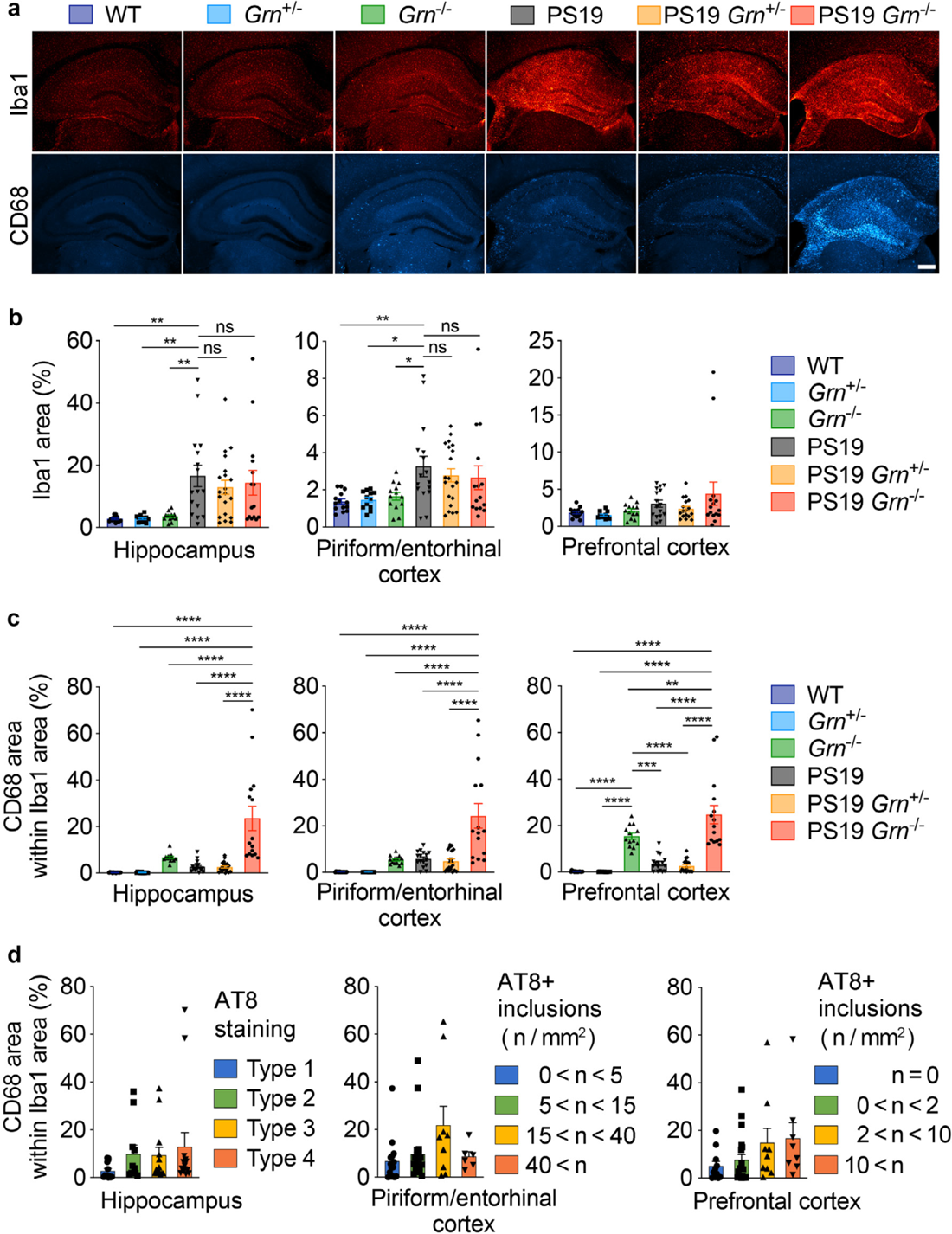
CD68-positive microglia are not associated with increased tau inclusions in PS19 mice with PGRN reduction. **a.** Representative images of Iba1 and CD68 double staining of the hippocampus of 9-12-month-old 6 genotypes. Bar, 300 μm. **b.** Quantification of Iba1 area and in 9-12-month-old 6 genotypes. Mean ± SEM, n = 12– 20/genotype, *p < 0.05, **p < 0.01; ****p < 0.0001; One-way ANOVA with Dunnett’s post hoc test. **c.** Quantification of CD68 area within Iba1 area (%) in 9-12-month-old 6 genotypes. Mean ± SEM, n = 12–20/genotype, **p < 0.01, ***p < 0.001; ****p < 0.0001; One-way ANOVA with Tukey post hoc test. **d.** No association of CD68 area within Iba1 area with AT8 staining patterns in the hippocampus or with the number of AT8+ inclusions in the piriform/entorhinal cortex or in the prefrontal cortex. n = 51 mice. Four groups in each graph are not significantly different by Kruskal-Wallist test.

### Tau phosphorylation and gliosis are not associated with increased tau inclusions in PS19 mice with PGRN reduction

We next sought to elucidate the mechanisms underlying increased tau inclusions in PS19 mice with PGRN reduction. We first examined the status of tau phosphorylation in the cortex of these animals. Immunoblot analysis using anti-total tau and phospho-tau antibodies (p-S199/S202, p-S202/T205, p-S356, and p-S396/S404) of the RAB and RIPA-soluble fractions revealed no significant changes in total tau and phospho-tau levels in the cortex of PS19 mice with PGRN reduction (Extended Data Fig. 4a-d).

Several recent studies have suggested an important role of microglial activation/neuroinflammation in driving tau pathology ^46, 51–53^. PGRN has also been implicated in microglial activation ^5, 6, 54^. Therefore, we next examined whether microgliosis is associated with increased tau inclusions in PS19 mice with PGRN reduction using microglial markers Iba1 and CD68. Immunohistochemistry showed an increase in Iba1-immunoreactivity in PS19 brains but no significant difference between PS19, PS19 *Grn*^+/−^ and PS19 *Grn*^−/−^ brains (Fig. 4a,b, and Extended Data Fig. 2b-e). Similar to our previous studies using the APP/PS1 cerebral A*β*-amyloidosis mouse model ^26^, CD68-immunoreactivity within Iba1 area was significantly increased in PS19 *Grn*^−/−^ mice. However, the increase was not observed in PS19 *Grn*^+/−^ mice (Fig. 4c). In addition, the CD68 levels were not significantly correlated with AT8 staining types in the hippocampus or with the number of tau inclusions in the piriform/entorhinal and prefrontal cortex in our cohorts (Fig. 4d). We also examined astrocytosis of the 6 genotypes using GFAP staining. Similar to the Iba1 staining, GFAP immunoreactivity was increased in PS19 brains, but it was not significantly affected by PGRN reduction (Extended Data Fig. 5). Thus, tau phosphorylation and glial marker changes were not associated with increased tau inclusions in PS19 mice with PGRN reduction.

**Fig. 5:**
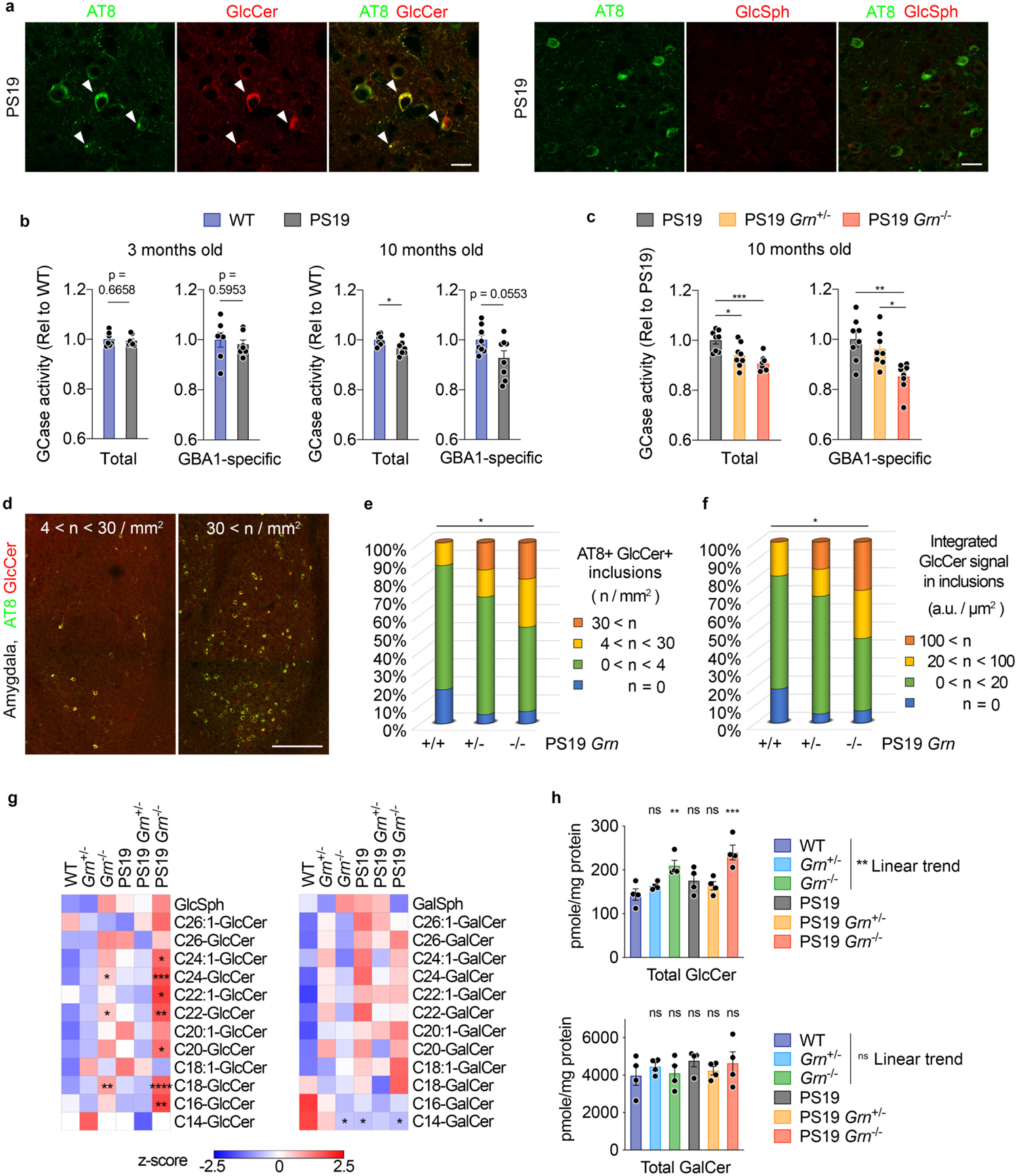
PGRN reduction attenuates GCase activity while tau inclusions are immunoreactive for GlcCer. **a.** Representative confocal images of AT8 and GlcCer or GlcSph co-staining in the amygdala region of PS19 mice. Bar, 20 μm. **b.** Total and GBA1-specific GCase activity of WT and PS19 cortices at 3 and 10 months of age. Mean ± SEM, n = 7 or 8 mice per genotype for 3 or 10 months of age, respectively. *p < 0.05; unpaired t-test. **c.** Total and GBA1-specific GCase activity of PS19, PS19 *Grn*^+/−^, and PS19 *Grn*^−/−^ cortices at 10 months of age. Mean ± SEM, n = 8 mice per genotype. *p < 0.05, **p < 0.01, ***p < 0.001; One-way ANOVA with Tukey’s post hoc test. **d.** Representative of confocal images of AT8 and GlcCer co-staining in the amygdala region. Bar, 200 μm. **e.** Distribution of the number of AT8 and GlcCer double-positive inclusions in the amygdala of 3 genotypes at 9-12 months of age. n = 15–20 mice per genotype, *p < 0.05; Chi-square test for trend (3 rows (PS19, PS19 *Grn*^+/−^, versus PS19 *Grn*^−/−^) and 2 columns (0 < n < 4 versus 4 < n /mm^2^)). **f.** Distribution of integrated GlcCer intensity in AT8 and GlcCer double-positive inclusions in the amygdala of 3 genotypes at 9-12 months of age. n = 15–20 mice per genotype, *p < 0.05; Chi-square test for trend (3 rows (PS19, PS19 *Grn*^+/−^, versus PS19 *Grn*^−/−^) and 2 columns (0 < n < 20 versus 20 < n /mm^2^)). **g.** Heatmaps showing levels of GlcCer and GalCer species in the cortex of 6 genotypes at 10 months of age. n = 4 mice per genotype. *p < 0.05, **p < 0.01, ***p < 0.001, ****p < 0.001; One-way ANOVA with Dunnett’s post hoc test comparing to WT. **h.** Total GlcCer or GalCer levels in the cortex of 6 genotypes at 10 months of age. Mean ± SEM, n = 4 mice per genotype. **p < 0.01, ***p < 0.001; One-way ANOVA with Dunnett’s post hoc test comparing to WT and One-way ANOVA with test for linear trend using only WT, *Grn*^+/−^, and *Grn*^−/−^.

### PGRN reduction attenuates GCase activity and increases GlcCer- and p-*α*-syn-positive inclusions in PS19 mice

We then considered whether PGRN reduction cell-autonomously affects tau inclusions in neurons. Co-staining with anti-PGRN and AT8 antibodies revealed no incorporation of PGRN into tau inclusions in PS19 mice (Extended Data Fig. 6a). Thus, it is unlikely that PGRN prevents formation of tau inclusions by directly binding to tau. Interestingly, we observed fewer PGRN-positive puncta in neurons with AT8-positive inclusions compared to ones without the inclusions (Extended Data Fig. 6a).

**Fig. 6:**
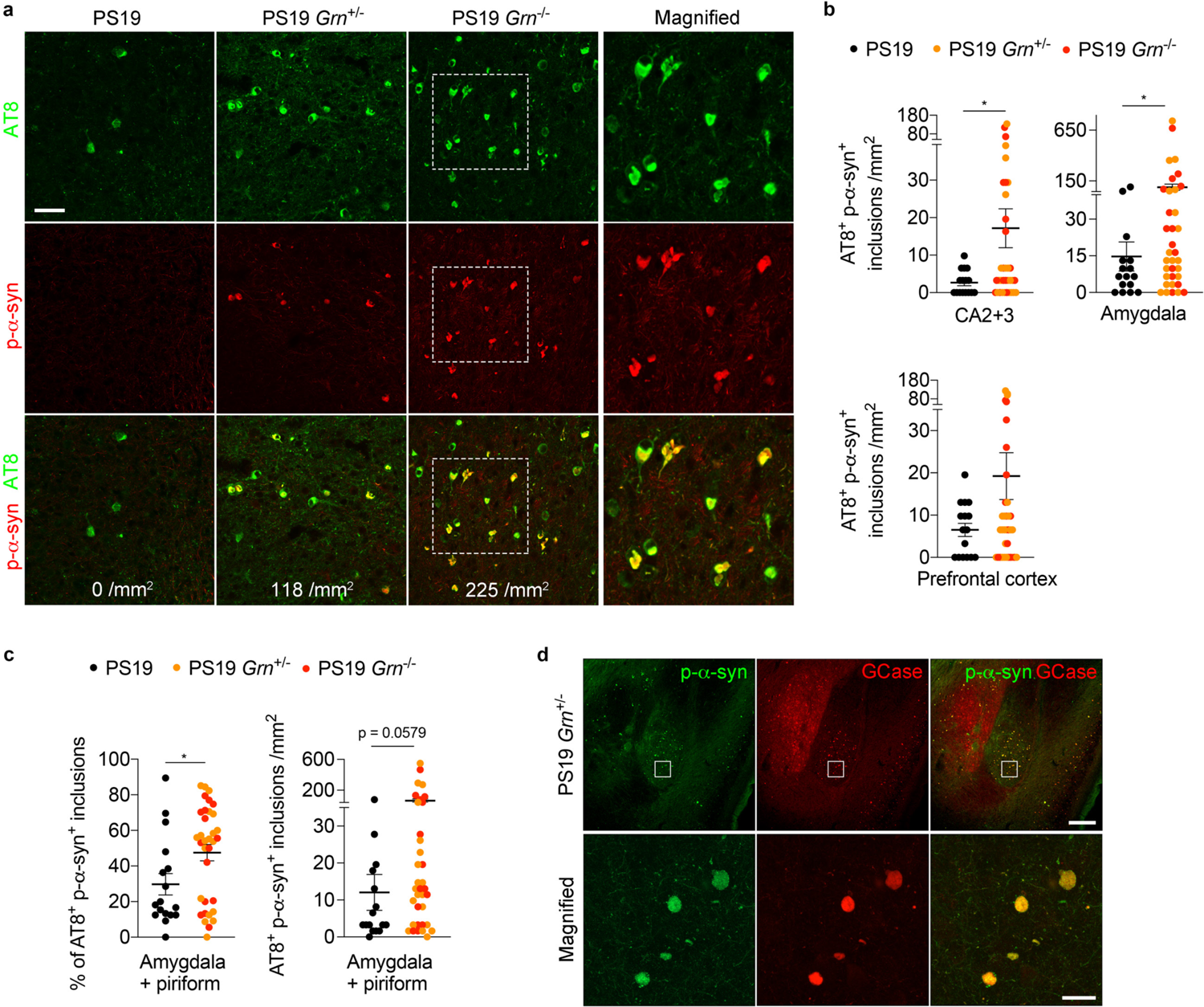
PGRN reduction exacerbates *α*-syn co-accumulation in PS19 mice. **a.** Representative confocal images of AT8 and p-*α*-syn co-staining in the amygdala region of PS19, PS19 *Grn*^+/−^ or PS19 *Grn*^−/−^ mice. Square dotted boxes indicate regions where magnified images are presented. The number of AT8 and p-*α*-syn double positive inclusions was shown on the bottom of each panel. Bar, 50 μm. **b.** Quantification of the number of AT8 and p-*α*-syn double positive inclusions of the CA2 and CA3, amygdala, and prefrontal cortex of 3 genotypes (PS19, PS19 *Grn*^+/−^, and PS19 *Grn*^−/−^ mice) at 9-12 months of age. Mean ± SEM, n = 15–20 mice per genotype, *p < 0.05; Mann-Whitney test. **c.** Quantification of the percentage (AT8^+^ p-*α*-syn^+^ inclusions per AT8^+^ inclusions) and number of AT8 and p-*α*-syn double positive inclusions in the amygdala plus piriform cortex region of PS19, PS19 *Grn*^+/−^, and PS19 *Grn*^−/−^ mice at 9-12 months of age. At least 7 AT8-positive inclusions were examined in each mouse. The averages of the number of AT8-positive inclusions examined per mouse are as follows: PS19 = 18.94 ± 4.285, PS19 *Grn*^+/−^ = 55.45 ± 22.63, PS19 *Grn*^−/−^ = 56.67 ± 23.20. Mean ± SEM, n = 15-20 mice per genotype. *p < 0.05; Mann-Whitney test. **d.** Representative confocal images of p-*α*-syn and GCase co-staining in the amygdala region of PS19 and PS19 *Grn*^+/−^ mice. Bar, 200 μm. The bottom panels show a high magnification of white square area in the top panels. Bar, 20 μm.

Recent studies have reported physical interaction between PGRN and *β*-glucocerebrosidase (GCase), a lysosomal enzyme that cleaves the glycosphingolipids glucosylceramide (GlcCer) and glucosylsphingosine (GlcSph). In addition, PGRN deficiency was shown to result in destabilization and/or reduced activity of GCase ^55–57^. Interestingly, mice with reduced GCase activity have been reported to develop tau pathology ^58–60^. We confirmed both the PGRN-GCase interaction using co-immunoprecipitation (co-IP) assay with HEK293T cells (Extended Data Fig. 5b,c) and the reduced GCase activity using 5-month-old *Grn*^−/−^ cortices (Extended Data Fig. 5d). Thus, we next examined whether GCase and GlcCer/Sph are involved in increased formation of tau inclusions upon PGRN reduction. Immunohistochemistry with anti-GlcCer and GlcSph antibodies revealed that tau inclusions in

PS19 mice were consistently immunoreactive for GlcCer but not GlcSph (Fig. 5a and Extended Data Fig. 5e,f), suggesting a potential involvement of GlcCer in tau inclusion formation. Importantly, specificity of the anti-GlcCer antibody has been previously demonstrated ^61, 62^ and the antibody has been commonly used for immunostaining ^63–66^. We also confirmed an increase in GlcCer and GlcSph immunoreactivity in the brains of mice treated with GCase inhibitor conduritol B epoxide (CBE) (Extended Data Fig. 6g-i). GCase activity assay using cortical lysates showed that while there was no difference in GCase activity between WT and PS19 brains at 3 months of age, total GCase activity was slightly but significantly decreased in PS19 brains at 10 months of age (Fig. 5b). In addition, total GCase and GBA1-specific activity of PS19 brains were further attenuated in both PS19 *Grn*^+/−^ and PS19 *Grn*^−/−^ brains (Fig. 5c). Consistent with the reduced GCase activity, immunohistochemistry showed gene dosage-dependent increases in GlcCer-positive tau inclusions and integrated GlcCer intensity within tau inclusions in PS19 mice with PGRN reduction (Fig. 5d-f). To determine levels of GlcCer and galactosylceramide (GalCer) species, we further performed SFC-MS/MS analysis using lipid extracts from cortical lysates of 6 genotypes. We found an increase in some of GlcCer species and total GlcCer levels in *Grn*^−/−^ and PS19 *Grn*^−/−^ cortices. In addition, there was a linear trend in the increase of total GlcCer levels between WT, *Grn*^+/−^ and *Grn*^−/−^ genotypes, although unexpectedly we did not observe the trend between PS19, PS19 *Grn*^+/−^ and PS19 *Grn*^−/−^ genotypes (Fig. 5g,h and Extended Data Fig. 7). Importantly, PGRN reduction did not cause any significant changes in GalCer species, which are also cerebrosides but are not substrates of GCase (Fig. 5g,h and Extended Data Fig. 7). These results suggest that PGRN reduction impairs GCase activity and that an increase in GlcCer accumulation may promote formation of tau inclusions in PS19 mice.

**Fig. 7:**
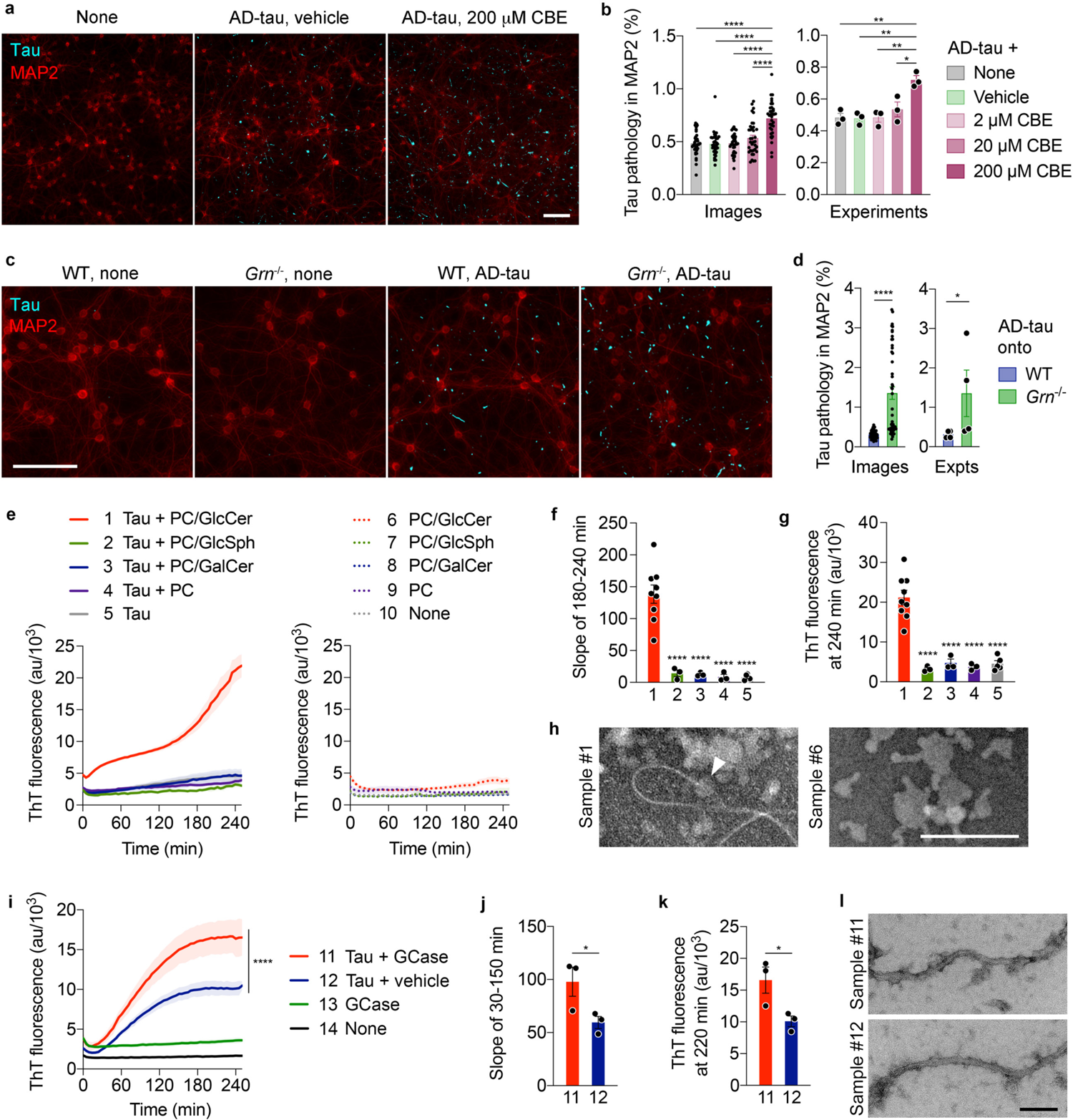
GlcCer and GCase promote tau aggregation *in vitro*. **a.** Representative images of mouse tau and MAP2 co-staining of WT neurons at DIV21 after AD-tau seeding in the presence of vehicle or 200 μM CBE. Bar, 100 μm. **b.** Quantification of endogenous mouse tau-positive area within MAP2 area. Neurons were seeded with AD-tau in the absence or presence of vehicle, 2 μM CBE, 20 μM CBE or 200 μM CBE. Experiments were performed in triplicate, and four images were unbiasedly taken from each well. Mean ± SEM, n = 36 images or 3 experiments, *p < 0.05, **p < 0.01, ****p < 0.0001; One-way ANOVA with Tukey’s post hoc test. **c.** Representative images of mouse tau and MAP2 co-staining of WT and *Grn*^−/−^ neurons at DIV21 after seeding AD-tau. Bar, 100 μm. **d.** Quantification of endogenous mouse tau-positive area within MAP2 area. Experiments were performed in triplicate, and four images were unbiasedly taken from each well. Mean ± SEM, n = 48 images or 4 experiments, *p < 0.05, ****p < 0.0001; Mann-Whitney test. **e.** ThT assay of heparin-induced aggregation of P301S tau in the presence or absence of PC, PC/GlcCer, PC/GlcSph, or PC/GalCer lipid dispersions. Heparin was included in all samples except for the sample #10. Mean ± SEM, n = 9 (for the sample #1 and 6), 5 (for the sample #5 and 10), or 3 (for the other samples) experiments, each preformed in triplicate. **f.** Slopes determined by linear regression from 180 to 240 min in **e**. ****p < 0.0001; One-way ANOVA with Tukey’s post hoc test. **g.** ThT fluorescence at 240 min in **e**. ****p < 0.0001; One-way ANOVA with Tukey’s post hoc test. **h.** Representative negative-stain EM images of the sample #1 and #6. An arrowhead indicates a tau fibril elongated from PC/GlcCer lipid dispersions. Bar, 200 nm. **i.** ThT assay of heparin-induced aggregation of P301S tau in the presence or absence of GCase. Instead of GCase, the same volume of 50 mM sodium citrate pH 5.5 was added as a vehicle control to the sample #12 and #14. Heparin was included in all samples except for the sample #14. Mean ± SEM, n = 3 experiments, each preformed in triplicate. ****p < 0.0001; nonlinear regression with a sigmoidal model followed by extra sum-of-squares F test. **j.** Slopes determined by linear regression from 30 to 150 min in **i**. *p < 0.05; paired t-test. **k.** ThT fluorescence at 220 min in **i.** *p < 0.05; paired t-test. **l.** Representative negative-stain EM images of sample #1 and #2. Bar, 100 nm.

Given the known prominent association of GCase activity with synucleinopathy ^36, 37^, we also examined *α*-syn pathology in 3 genotypes (PS19, PS19 *Grn*^+/−^ and PS19 *Grn*^−/−^) using anti-phospho-Ser129 (p-*α*-syn) antibody. We observed a sparse p-*α*-syn immunoreactivity that is colocalized with AT8-positive inclusions in PS19 brains, which was significantly increased in the CA2, CA3 and amygdala regions upon PGRN reduction when a dominant model was tested (Fig. 6a,b and Extended Data Fig. 8a,c,d). The density of AT8 and p-*α*-syn double positive inclusions was also increased in PS19 mice with PGRN reduction when a large number of AT8-positive inclusions was analyzed using both the amygdala and piriform cortex regions (Fig. 6c). In addition, using immunohistochemistry with anti-GCase antibody we found a significant incorporation of GCase in the AT8 and p-*α*-syn double positive inclusions in PS19 mice with PGRN reduction (Fig. 6d and Extended Data Fig. 8b), while the co-aggregates were not immunoreactive for TDP-43 (Extended Data Fig. 8e). Thus, PGRN reduction also exacerbated *α*-synuclein and GCase but not TDP-43 co-accumulation in PS19 mice.

**Fig. 8:**
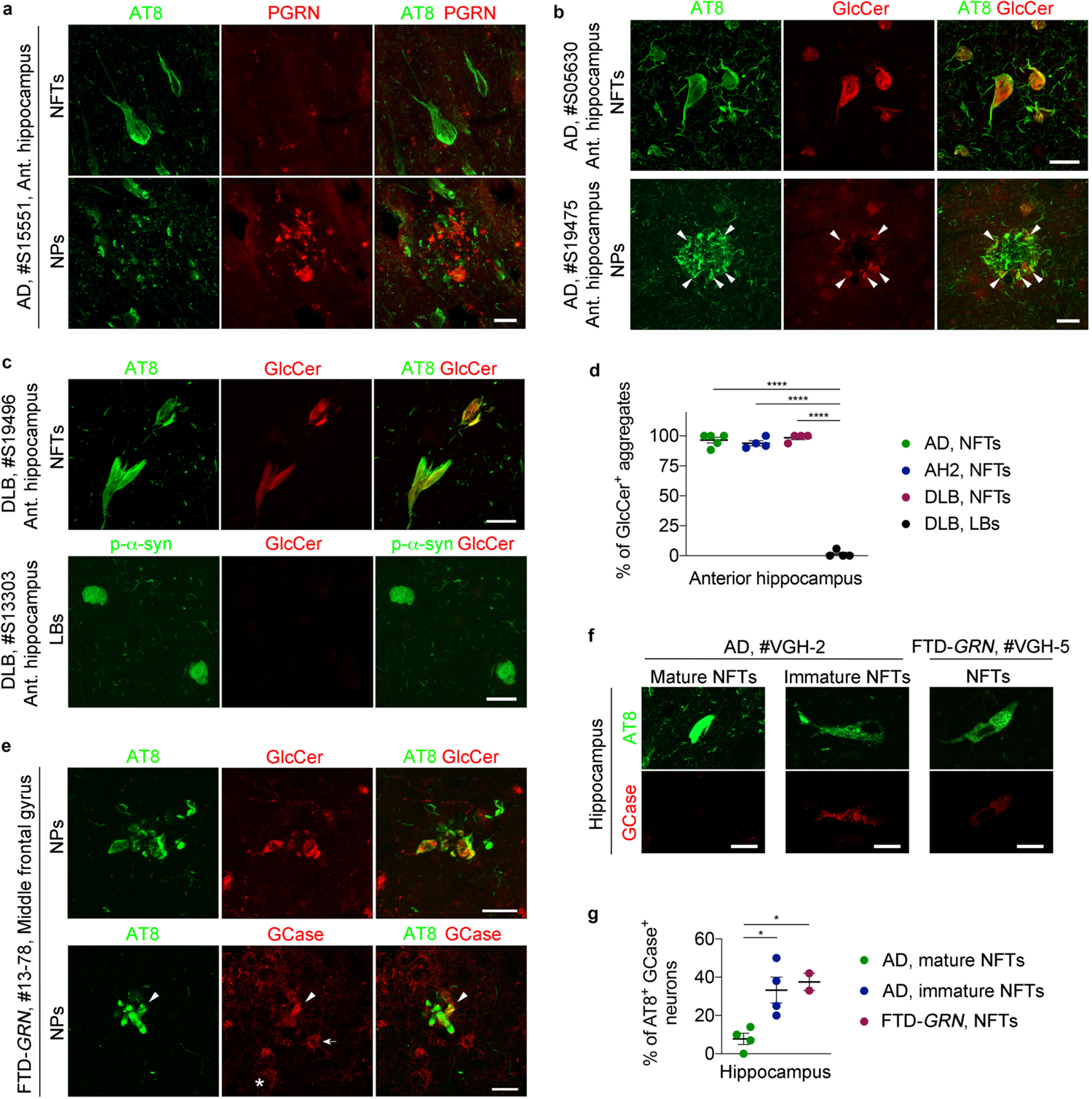
Accumulation of GlcCer and GCase to tau aggregates in AD, DLB, and FTD-*GRN* patients. **a.** Representative confocal images of AT8 and PGRN co-staining of NFTs and NPs in the anterior hippocampus region of AD brain. Bar, 20 μm. **b.** Representative confocal images of AT8 and GlcCer co-staining of NFTs and NPs in the anterior hippocampus region of AD brains. Arrowheads indicate GlCer labeling of some of NPs. Bar, 20 μm. **c.** Representative confocal images of AT8 and GlcCer co-staining of NFTs and LBs in the anterior hippocampus region of DLB brains. Bar, 20 μm. **d.** Quantification of percentage of GlcCer+ aggregates of NFTs of AD, AH2, DLB brains and LBs of DLB brain tissue. At least 10 NFTs were examined in each sample except for two DLB samples due to the low numbers of NFTs (2 and 6 NFTs). The mean number of NFTs examined in each group is as follows: AD = 15 ± 1.924, AH2 = 13.75 ± 1.652, DLB = 8.5 ± 2.986. At least 11 LBs were examined for each DLB brain and the mean number is 14 ± 1.472. Mean ± SEM, n = 4-5 brains, ****p < 0.0001; One-way ANOVA with Tukey’s post hoc test. **e.** Representative confocal images of AT8 and GlcCer or GCase of co-staining of NPs in the middle frontal gyrus of a FTD-*GRN* brain. Bar, 20 μm. An arrowhead indicates co-localization of GCase with AT8-positive neuritic plaques. An arrow indicates glial localization of GCase. An asterisk indicates lysosomal GCase in neurons. Bar, 20 μm. **f.** Representative confocal images of AT8 and GCase of co-staining of NFTs in the hippocampus of AD and FTD-*GRN* brain. Bar, 20 μm. **g.** Quantification of percentage of AT8+ GlcCer+ aggregates of NFTs in the hippocampus of AD and FTD-*GRN* brain. NFTs of AD were divided into two groups: mature and immature NFTs, based on their morphology. At least 48 NFTs for each group were examined in each sample except for one FTD-*GRN* sample due to the low number of NFTs (15 NFTs). The mean number of NFTs examined in each group is as follows: AD, mature NFTs = 56.25 ± 3.75, AH2, immature NFTs = 58.00 ± 4.673, FTD-*GRN* NFTs = 35.5 ± 20.50. Mean ± SEM, n = 2-4 patients, *p < 0.05; One-way ANOVA with Tukey’s post hoc test. See also Figure S6 and S7.

### GlcCer and GCase promotes tau aggregation *in vitro*

To further investigate the role of GCase and GlcCer in tau aggregation, we examined the effect of CBE in the well-established tau seeding assay using primary cultured neurons ^67, 68^. Tau fibrils extracted from AD brains (AD-tau) were seeded onto neurons with CBE or vehicle treatment and incubated for two weeks. Immunofluorescence using anti-mouse tau antibody revealed that a high concentration of CBE significantly promotes tau aggregation induced by AD-tau (Fig. 7a,b). Similarly, we also found that PGRN deficiency increases tau aggregation induced by AD-tau in neurons although no tau aggregation was detected in *Grn*^−/−^ neurons without AD-tau seeding (Fig. 7c,d). Interestingly, we also observed an increase in AT8 mean intensity in the piriform cortex of WT mice treated with CBE for 21 days although the inclusions were not detected in the mice likely due to the short period of time (Extended Data Fig. 6j). To examine whether accumulation of GlcCer by GCase inhibition directly affects tau aggregation, we next performed *in vitro* Thioflavin T (ThT) assay using recombinant P301S tau and lipid dispersions made of purified phosphatidylcholine (PC) with or without GlcCer, GlcSph, or GalCer. Surprisingly, we found that heparin-induced aggregation of P301S tau was dramatically accelerated in the presence of PC/GlcCer lipid dispersions, but not the other lipid dispersions, while the lipid dispersions have no effects on ThT fluorescence (Fig. 7e-g). Negative-stain EM analysis showed the presence of tau fibrils that appear to be elongated from the PC/GlcCer lipid dispersions (Fig. 7h). Together, these results suggest a possibility that GlcCer directly accelerates tau aggregation.

Given the incorporation of GCase in tau inclusions in PS19 mice with PGRN reduction, we also examined whether GCase affects tau aggregation *in vitro*. Similar to GlcCer, *in vitro* ThT assay revealed that recombinant GCase significantly accelerates heparin-induced P301S tau aggregation, while GCase alone did not spontaneously aggregate at the concentration we tested (Fig. 7i-k). We confirmed that BSA has no effects on the tau aggregation, excluding a possibility that total protein concentration modulates tau aggregation in this *in vitro* experimental setting (Extended Data Fig. 9). Negative-stain EM analysis showed that there is no significant morphological difference between fibrils with and without GCase (Fig. 7l). These results suggest that GCase protein itself could also promote tau aggregation.

### NFTs in human tauopathy brains are immunoreactive for GlcCer

Finally, we investigated the relationship between tau, PGRN, GCase, and *α*-syn in human autopsy brain with tauopathy. We first examined PGRN expression in tau pathology of AD. Similar to the results obtained from PS19 mice, we found no and very weak PGRN immunoreactivity in AT8-positive neurofibrillary tangles (NFTs) and neuritic plaques (NPs) of AD brains, respectively, even though PGRN signal was detected in neurons and significantly increased in dystrophic neurites near amyloid plaques (Fig. 8a and Extended Data Fig. 10b), which is partly consistent with a recent publication ^69^.

We then examined whether similar co-aggregation of GlcCer and GCase found in PS19 mice with PGRN reduction can be also observed in tau pathologies in anterior hippocampal and middle frontal gyrus sections from 5 AD brains, 4 with moderate AD pathological changes (AH2), and 1 frontotemporal dementia (FTD)-*GRN*. Strikingly, we detected high GlcCer positivity in NFTs of the anterior hippocampus in all of five AD brains and four AH2 brains we tested (Fig. 8b,d and Extended Data Fig. 10c). In addition, GlcCer signal was partially colocalized with NPs in AD brains (Fig. 8b and Extended Data Fig. 10d). The GlcCer immunoreactivty was also detected in NFTs of DLB brains, while Lewy bodies of the DLB brains were largely negative for GlcCer (Fig. 8c,d). Furthermore, we observed a partial colocalization of GlcCer and GCase with NPs in the middle frontal gyrus of an FTD-*GRN* brain with AD pathology (Fig. 8e). We further examined GCase and p-*α*-syn immunoreactivities using another set of paraffin-embedded hippocampal sections of 4 AD and 2 FTD-*GRN* brains. While mature NFTs showed low GCase positivity, immature NFTs and NFTs found in FTD-*GRN* exhibited higher GCase positivity (Fig. 8f.g). In 2 AD brains, we found dot-like aggregates of p-*α*-syn in a small subset of AT8-positive NFTs (Extended Data Fig. 10e). Similar dot-like aggregates of p-*α*-syn were also observed in our PS19 Grn cohort (Extended Data Fig. 8c,d). Of the AT8 and p-*α*-syn positive neurons, more than 60% showed GCase immunoreactivity (Extended Data Fig. 10f,g). Together, these results suggest GlcCer involvement and potential co-aggregation of GCase and/or *α*-syn in human tauopathy.

## Discussion

### The role for a lysosomal PGRN-GCase complex in tau aggregation

A striking observation of the present study is that tau inclusions in PS19 neurons were consistently immunoreactive for the glycosphingolipid GlcCer, a substrate of GCase. This was somewhat unexpected because GCase has been previously associated with synucleinopathy^36, 37^ rather than tauopathy. However, there are also several recent studies showing concomitant tau pathology in the brains of GCase mutant mice and Gaucher disease patients in which GCase is mutated ^58–60, 70^. Importantly, in the present study, NFTs and some of NPs of human AD, DLB, and FTLD-*GRN* brains were also GlcCer-immunoreactive. Consistent with our observation, previous studies have revealed the presence of glycolipids or cerebrosides in paired helical filaments of AD ^71, 72^. In neuronal culture, we showed that GCase inhibition exacerbates tau aggregation induced by AD-tau. *In vitro*, PC/GlcCer lipid dispersions dramatically promoted heparin-induced aggregation of P301S tau. Together, these results clearly suggest a critical role of GlcCer in tau aggregation. In future studies, it will be interesting to examine the prevalence of tau pathology in a large cohort of human subjects with GCase mutations.

We then found that both complete loss and haploinsufficiency of PGRN impair GCase activity while increasing GlcCer-positive tau inclusions in PS19 mice. These results suggest that a lysosomal PGRN-GCase complex regulates formation of tau inclusions by regulating accumulation of GlcCer. Previously, PGRN has been known to bind to GCase, but the roles in health and disease were not clear. The present study for the first time links a lysosomal PGRN-GCase complex to tauopathy. GCase protein itself was also incorporated into tau inclusions in PS19 mice with PGRN reduction. *In vitro,* similar to GlcCer, recombinant GCase protein promoted heparin-induced aggregation of P301S tau. PGRN deficiency reportedly causes insolubility or aggregation of GCase in both humans and rodents ^56, 57^. Thus, PGRN binding to GCase may also be crucial to stabilize GCase protein thereby preventing its co-aggregation with tau. Recent studies have shown a role for microglial activation in driving tau pathology ^46, 51–53^, and PGRN plays an important role in microglial activation ^5,6, 54^. However, in the present study, the extent of CD68+ microgliosis was not associated with either AT8 staining types or the number of tau inclusions. It is thus unlikely that PGRN reduction accelerates formation of tau inclusions via microglial activation.

Interestingly, we observed that neurons with tau inclusions exhibit little or no PGRN expression in both PS19 mice and human AD brains. Consistent with a previous study ^69^, PGRN immunoreactivity was absent in NFTs. As previously reported ^8, 69, 73^, our immunohistochemistry showed that PGRN was highly expressed in dystrophic neurites near amyloid plaques in AD. However, the PGRN-positive dystrophic neurites were poorly colocalized with AT8-positive NPs. We have found that GCase activity is a slightly but significantly attenuated in the cortex of PS19 mice at 10 months of age. Thus, there may be a feedback mechanism by which pathological tau affects PGRN expression and GCase activity to exacerbate its accumulation. Further investigation will be required to explore this possibility.

### Implications of increased *α*-syn co-pathology in PS19 mice with PGRN reduction

In the present study, PGRN reduction also increased *α*-syn co-accumulation in PS19 mice. Concomitant proteinopathies are frequently observed in aged brain with neurodegenerative diseases and influence their clinical course ^2–4^, suggesting that therapies targeting pathways that affect co-pathologies may be more effective than monotherapies focusing on a single disease-associated protein. Our results in the present study suggest that PGRN reduction may cause tau and *α*-syn accumulation by impairing GCase activity and increasing GlcCer levels in FTLD-*GRN* and in other neurodegenerative diseases, consistent with a large body of literatures showing an association between GCase and synucleinopathy^36, 37^. This also suggests that a lysosomal PGRN–GCase complex may be a common therapeutic target for comorbid proteinopathies in neurodegenerative disease. Similar to the effects of GlcCer on tau in the present study, a previous study has shown direct effects of GlcCer on *α*-syn aggregation ^63^. Destabilization and aggregation of GCase by PGRN reduction may also contribute to the tau and *α*-syn co-accumulation, since incorporation of GCase into Lewy bodies is reported in patients with parkinsonism ^37^.

### Differential effects of PGRN deficiency on symptoms of tauopathy

In the present study, our results revealed a differential role of PGRN in memory and disinhibited behaviors in the setting of tauopathy. PGRN reduction worsens disinhibited behaviors while improving a spatial memory deficit in PS19 mice. The improvement was coupled with attenuated neurodegeneration and transcriptomic changes in the hippocampus. PS19 mice, a tauopathy model expressing human P301S tau under the control of the mouse prion protein promoter, display a wide range of behavioral deficits observed in human tauopathies such as AD and FTLD ^40, 41^. In FTD patients, memory function is preserved until late stage of the disease, while disinhibition is a key clinical symptom of behavioral variant FTD patients ^74^. Similarly, in the presence of tau pathology, reduction of PGRN, a FTD-associated protein, might attenuate tau-driven memory impairment while exacerbating FTD-like disinhibited behavior in PS19 mice. Intriguingly, in PS19 mice with PGRN reduction, the rescue of memory impairment and neurodegeneration was accompanied by an increase in tau inclusions. This finding seems somewhat contradictory but is consistent with recent studies using PS19 mice. Type 2 AT8 staining pattern, which is characterized by frequent tangle-like staining, has been shown to correlate with less brain atrophy, while type 4, strong but diffuse staining without large somatic inclusions, was frequently observed in brains with severe atrophy ^43^. Another published study showed a rescue of neurodegeneration and memory deficits in parallel with an increase in NFTs ^75^. Thus, formation of somatic tau inclusions might protect against hippocampal neurodegeneration and functions, although it is important to validate the protective role of tau inclusions using different approach and tauopathy models.

How PGRN reduction exacerbates disinhibited behaviors in tauopathy is less clear. In the present study, we also observed a trend toward an increase in co-accumulation of tau and *α*-syn in the prefrontal cortex, a brain region implicated in executive functions including inhibitory control ^76^. An increase in co-pathology of tau and *α*-syn might perturb neuronal networks regulating inhibitory control in the prefrontal cortex despite rescue of neurodegeneration in the hippocampus.

The differential effects on symptoms have a significant implication for PGRN’s role in AD. While improvement of memory in PS19 mice with PGRN reduction might not explain the mechanism of increased AD risk by *GRN* rs5848 T allele, exacerbation of disinhibited behaviors in combination with increased tau inclusions could do. It will be interesting to examine whether similar alteration of symptoms is observed in AD patients with the *GRN* rs5848 T allele.

### The effects of PGRN haploinsufficiency in PS19 mice

The present study shows clear effects of PGRN haploinsufficiency in PS19 mice, closely recapitulating those of the *GRN* mutations or rs5848 variant in AD. This is a striking contrast to most, but not all, previous studies describing no obvious effects of PGRN haploinsufficiency in mice ^5, 6^. Attenuated behavioral deficits and neurodegeneration, increase in tau and p-*α*-syn inclusions, reduced GCase activity were all observed not only in PS19 *Grn*^−/−^ mice but also in PS19 *Grn*^+/−^ mice. As described above, our data suggest that phenotypes observed in the present study are due, at least in part, to reduced GCase activity and stability in PS19 mice with PGRN reduction. Thus, direct and primary effects of loss of lysosomal PGRN-GCase complex after PGRN reduction could explain obvious phenotypes in PS19 *Grn*^+/−^ mice. This raises an intriguing possibility that phenotypes due to reduced GCase activity or levels might be observed even in *Grn* heterozygous mice, although it also requires further investigation.

### Transcriptomic analysis of PGRN deficient mice

Bulk RNA-seq/transcriptomic analysis using PGRN deficient mice from our previous study and others has shown up-regulation of lysosomal genes and microglial TYROBP gene network including C1q genes even at early stage of mouse lifetime ^39, 77, 78^. Our snRNA-seq detected only limited changes in such genes for *Grn*^−/−^ nuclei without the PS19 transgene in comparison to WT. The statistically significant up-regulation of lysosomal genes in *Grn*^−/−^ mice was in most instances less than two-fold. We did confirm lysosomal and microglial changes (e.g., DPPII and CD68) at protein levels in the hippocampus of *Grn*^−/−^ mice by immunohistochemistry (Fig. 4, DPPII data not shown), consistent with our previous studies ^26, 39^. The less prominent transcriptomic changes for these pathways may relate to the significant pools of RNA outside the nucleus being discarded during nuclei isolation, similar to a recent study on a different topic^79^.

## Conclusion

The present study revealed an unexpected role of glycosphingolipid GlcCer in tau aggregation and demonstrated that a lysosomal PGRN-GCase complex titrates accumulation of GlcCer to regulate formation of tau and *α*-syn inclusions, which alters symptoms and neurodegeneration in tauopathy (Extended Data Fig. 10a). Our findings provide novel insights into roles lysosomes play in proteinopathy and advance our understanding of PGRN as a risk factor for AD and PD. It is likely that the PGRN-GCase complex plays an important role not only in Gaucher disease and FTLD but also in other neurodegenerative proteinopathies.

## Author Contributions

Conceptualization, H.T. and S.M.S.; Methodology, H.T., S.H.N., M.T.C., G.W., and S.M.S.; Investigation, H.T., S.B., S.H.N., M.T.C., and G.W; Writing – Original Draft, H.T. and S.M.S.; Writing – Review & Editing, all; Funding Acquisition, S.M.S.; Resources, I.M. and S.M.S; Supervision, S.M.S.

## Acknowledgments

We thank Marc Llaguno and Kimberley Gibson of the Yale School of Medicine Electron Microscopy Core Facility for expert technical assistance. We are grateful to the Banner Sun Health Research Institute Brain and Body Donation Program of Sun City, Arizona for the provision of human brain tissues. The Brain and Body Donation Program has been supported by the National Institute of Neurological Disorders and Stroke (U24 NS072026 National Brain and Tissue Resource for Parkinson’s Disease and Related Disorders), the National Institute on Aging (P30 AG19610 Arizona Alzheimer’s Disease Core Center), the Arizona Department of Health Services (contract 211002, Arizona Alzheimer’s Research Center), the Arizona Biomedical Research Commission (contracts 4001, 0011, 05-901 and 1001 to the Arizona Parkinson’s Disease Consortium) and the Michael J. Fox Foundation for Parkinson’s Research. Human brain tissues were also obtained from the NIH NeuroBioBank. S.H.N. received a PhD fellowship from Boehringer Ingelheim Fonds. This work was supported in part by the Lipidomics Shared Resource, Hollings Cancer Center, Medical University of South Carolina (P30 CA138313 and P30 GM103339). This work was supported by grants from the Falk Medical Research Trust and from the N.I.H. to S.M.S.

## Extended data

**Extended Data Fig. 1:**
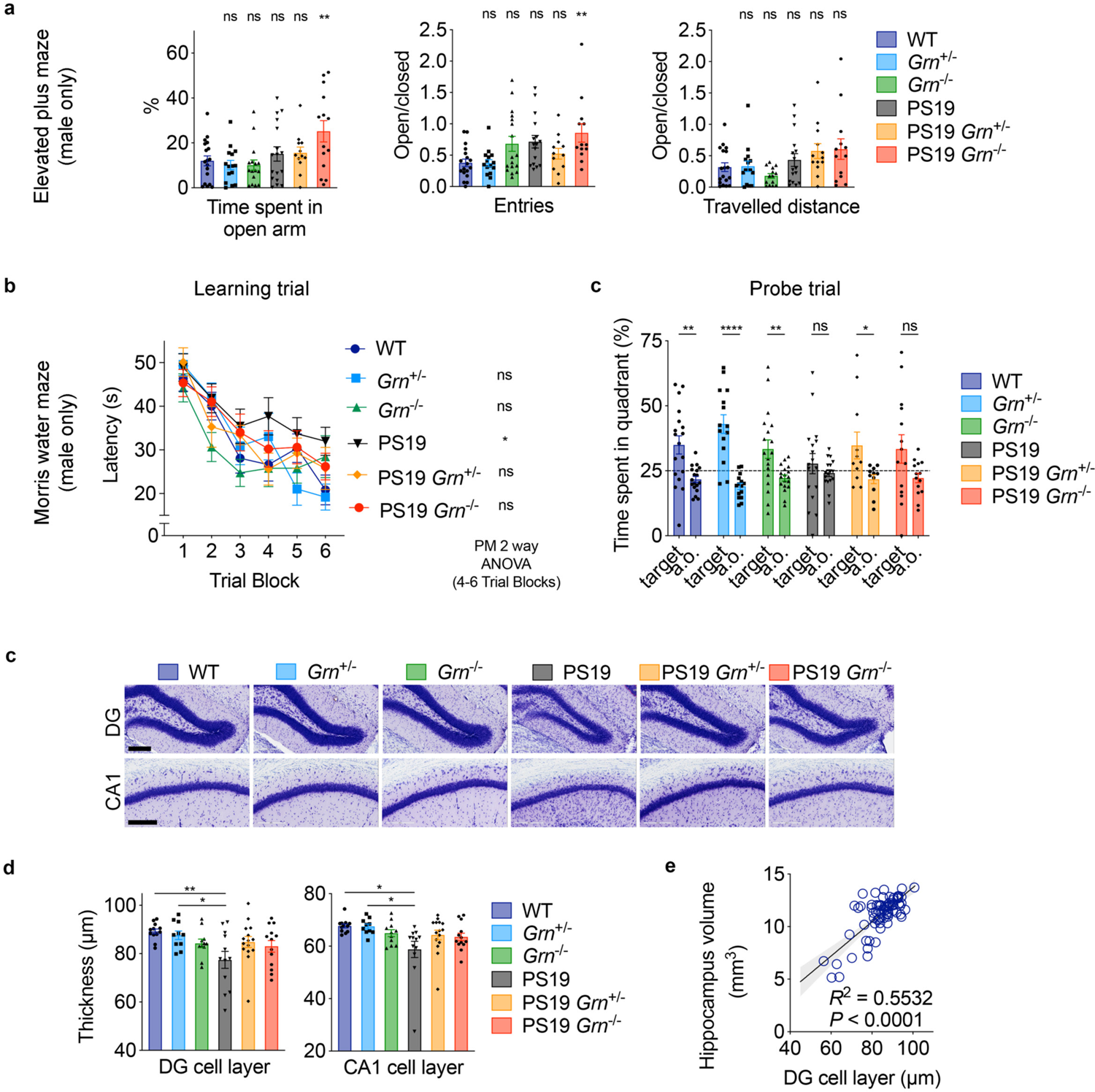
PGRN reduction exacerbates disinhibition while improving a memory deficit in male PS19 mice. **a.** Elevated plus maze (EPM) of male 6 genotypes (WT, *Grn*^+/−^, *Grn*^−/−^, PS19, PS19 *Grn*^+/−^, and PS19 *Grn*^−/−^ mice) at 10-11 months of age showing time spent in the open arms (%), and ratio of number of entries into the open versus closed arms ration of distance traveled in the open versus closed arms of the maze. Mean ± SEM, n = 11–18 mice per genotype, **p < 0.01; one-way ANOVA with Dunnett’s post hoc test. **b.** Morris water maze (MWM) learning trial of male 6 genotypes at 10-11 months of age. Spatial learning is plotted as latency to find hidden platform. Mean ± SEM, n = 11–18 mice per genotype *p < 0.05; repeated measures two-way ANOVA with Dunnett’s post hoc test using 4–6 Trial Blocks. There was also interaction between Trial Block and genotype (p = 0.0049). **c.** MWM probe trial of male 6 genotypes at 10-11 months of age. Percentage of time spent in the target quadrant and averaged time spent in all other (a.o.) quadrants. Mean ± SEM, n = 11–18 mice per genotype *p < 0.05, **p < 0.01, ****p < 0.0001; t test with Welch’s correction. **d.** Representative images of Nissl staining of the dentate gyrus (DG) and CA1 region of hippocampus from 6 genotypes. Bar, 200 μm. **e.** Thickness of the DG granule cell layer and CA1 pyramidal neuronal layer in 6 genotypes at 9-11 months of age. Mean ± SEM, n = 10–15 mice per genotype, *p < 0.05; **p < 0.01; One-way ANOVA with Tukey’s post hoc test. **f.** Correlation between DG granule cell layer thickness and hippocampal volume. n = 73 mice, R^2^ = 0.5532, p < 0.0001; Pearson correlation analysis.

**Extended Data Fig. 2:**
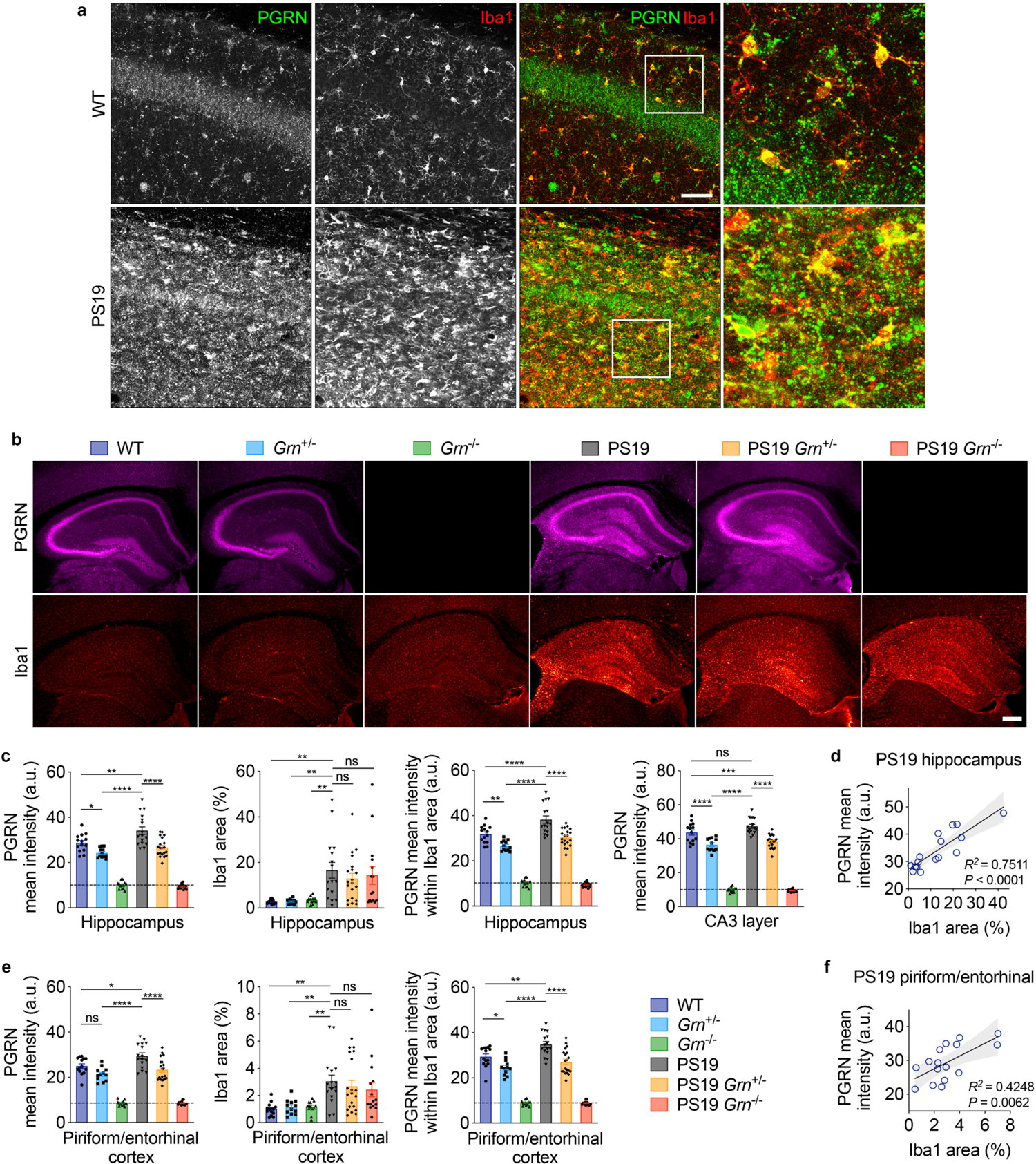
PGRN is increased in PS19 mice mainly due to microgliosis and is decreased in PS19 *Grn*^+/−^ mice. **a.** Representative confocal images of PGRN and Iba1 co-staining of the C1A region of hippocampus of 11-month-old WT and PS19 mice. Bar, 50 μm. **b.** Representative images of PGRN and Iba1 co-staining of the hippocampus of 6 genotypes (WT, *Grn*^+/−^, *Grn*^−/−^, PS19, PS19 *Grn*^+/−^, and PS19 *Grn*^−/−^ mice) at 9-12 months of age. Bar, 300 μm. **c.** Quantification of PGRN mean intensity, Iba1 area (%), and PGRN mean intensity within Iba1 area PGRN mean intensity of CA2 and CA3 neuronal layer in the hippocampus of 9-12-month-old 6 genotypes. Mean ± SEM, n = 12–20 mice per genotype, *p < 0.05, **p < 0.01; ***p < 0.001; ****p < 0.0001; One-way ANOVA with Tukey or Dunnett’s post hoc test (for Iba1 area). **d.** Correlation between Iba1 area and PGRN mean intensity in the hippocampus of PS19 mice. n = 16 mice, R^2^ = 0.7511, p < 0.0001; Pearson correlation analysis **e.** Quantification of PGRN mean intensity, Iba1 area (%), PGRN mean intensity within Iba1 area in the piriform/entorhinal cotex of 9-12-month-old 6 genotypes. Mean ± SEM, n = 12–20 mice per genotype, *p < 0.05, **p < 0.01; ****p < 0.0001; One-way ANOVA with Tukey or Dunnett’s post hoc test (for Iba1 area). **f.** Correlation between Iba1 area and PGRN mean intensity in the piriform/entorhinal cortex of PS19 mice. n = 16 mice, R^2^ = 0.4248, p < 0.0062; Pearson correlation analysis.

**Extended Data Fig. 3:**
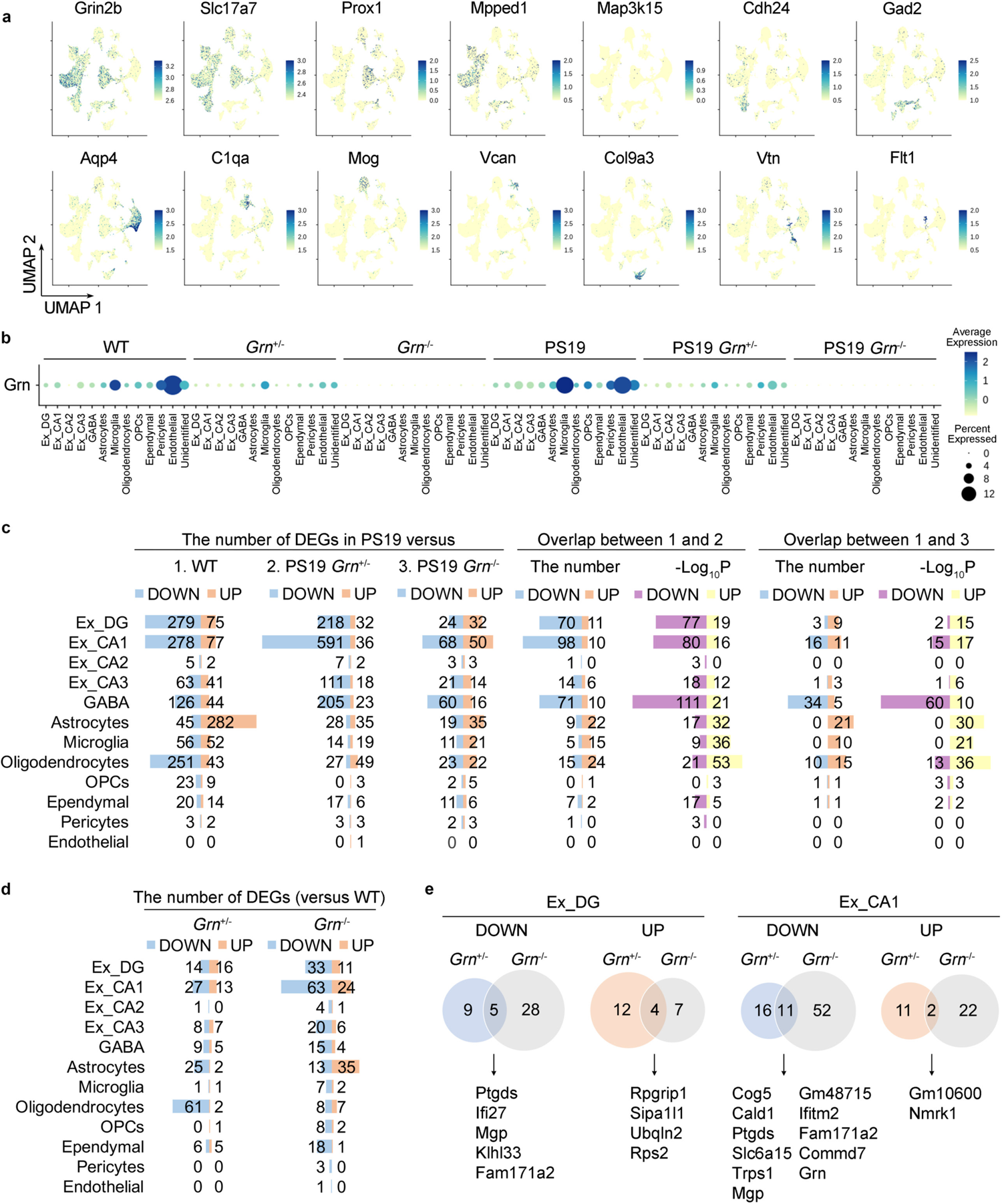
Marker gene expression and differential gene expression analyses in *Grn*^+/−^ or *Grn*^−/−^ vs. WT. **a.** UMAP plot colored by expression levels of marker genes: Grin2b (pan neurons), Slc17a7 (excitatory neurons), Prox1 (DG granule cells), Mpped1 (CA1 pyramidal cells), Map3k15 (CA2 pyramidal cells), Cdh24 (CA3 pyramidal cells), Gad2 (GABAergic neurons), Aqp4 (astrocytes), C1qa (microglia), Mog (oligodendrocytes), Vcan (oligodendrocytes precursor cells (OPCs)), Col9a3 (ependymal), Vtn (pericytes), Flt1 (endothelial). **b.** Dot plot showing expression of Grn in each cell type of 6 genotypes. **c.** Bar graphs showing the number of differentially-expressed genes (DEGs) (defined by Wilcoxon-rank-sum test, adjusted p < 0.01, |log(FC)| > 0.25) in PS19 nuclei compared to WT, PS19 *Grn*^+/−^, or PS19 *Grn*^−/−^ nuclei, and the number of overlapping genes between DEGs from PS19 vs. WT and from PS19 vs. PS19 *Grn*^+/−^ or from PS19 vs. PS19 *Grn*^−/−^. Overlapping p-values (-log_10_P) by Fisher’s exact test of each overlap are also bar-graphed. **d.** Bar graphs showing the number of differentially-expressed genes (DEGs) in *Grn*^+/−^ or *Grn*^−/−^ nuclei compared to WT nuclei (Wilcoxon-rank-sum test, adjusted p < 0.01, |log(FC)| > 0.25). **e.** Overlaps of DEGs in Ex_DG and Ex_CA1 between *Grn*^+/−^ and *Grn*^−/−^ nuclei.

**Extended Data Fig. 4:**
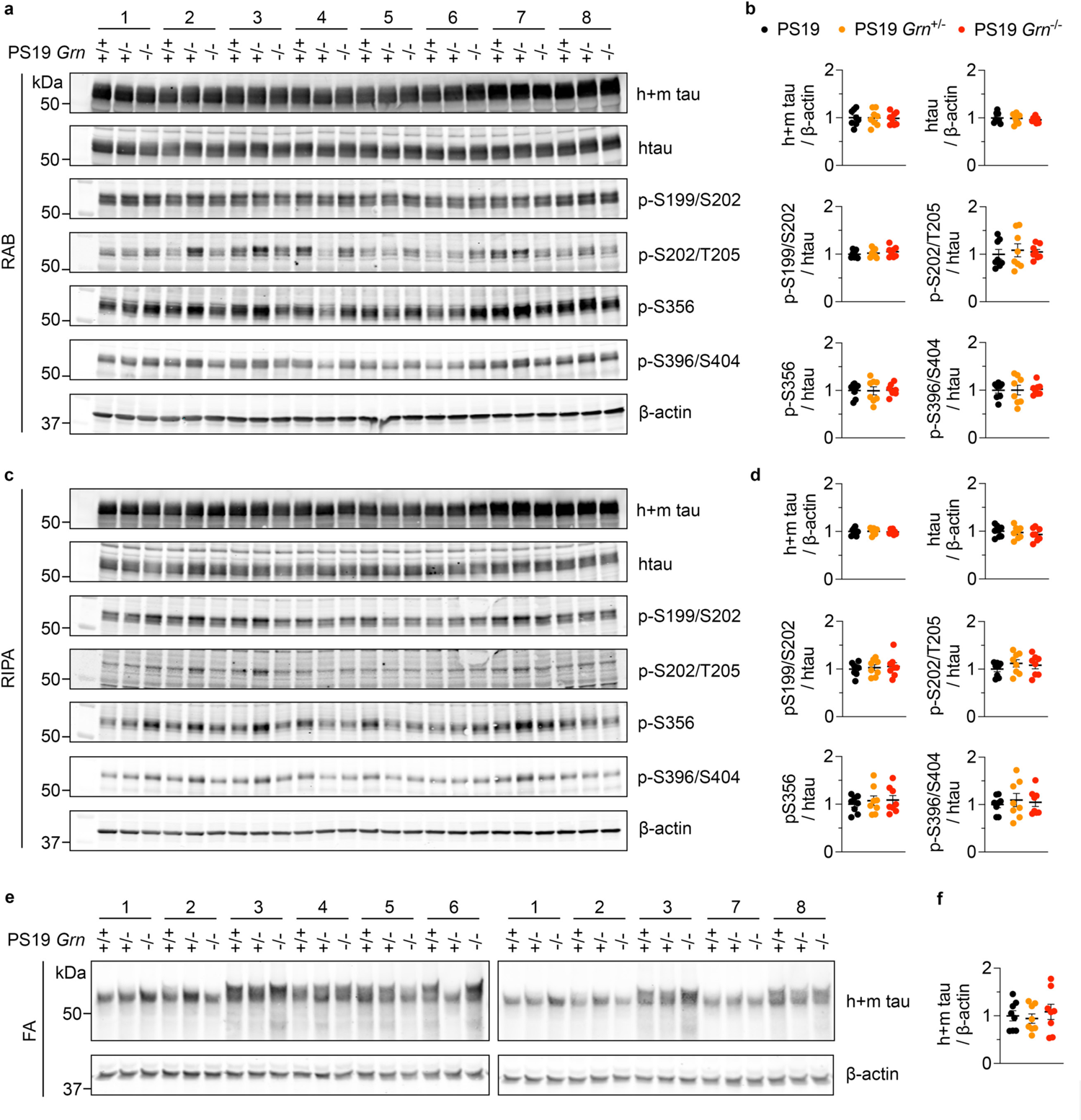
PGRN deficiency has no significant effects on tau phosphorylation in the RAB and RIPA-soluble fractions of PS19 mice. **a.** Immunoblot using anti-total tau, phospho-tau, and *β*-actin antibodies of the RAB-soluble fraction of 3 genotypes at 10 months of age. **b.** Quantification of immunoblot in **a**. Mean ± SEM, n = 8 mice per genotype. **c.** Immunoblot using anti-total tau, phospho-tau, and *β*-actin antibodies of the RIPA-soluble fraction of 3 genotypes at 10 months of age. **d.** Quantification of immunoblot in **c**. Mean ± SEM, n = 8 mice per genotype. **e.** Immunoblot using anti-total tau and *β*-actin antibodies of the FA-soluble fraction of 3 genotypes at 10 months of age. **f.** Quantification of immunoblot in **e**. Mean ± SEM, n = 8 mice per genotype.

**Extended Data Fig. 5:**
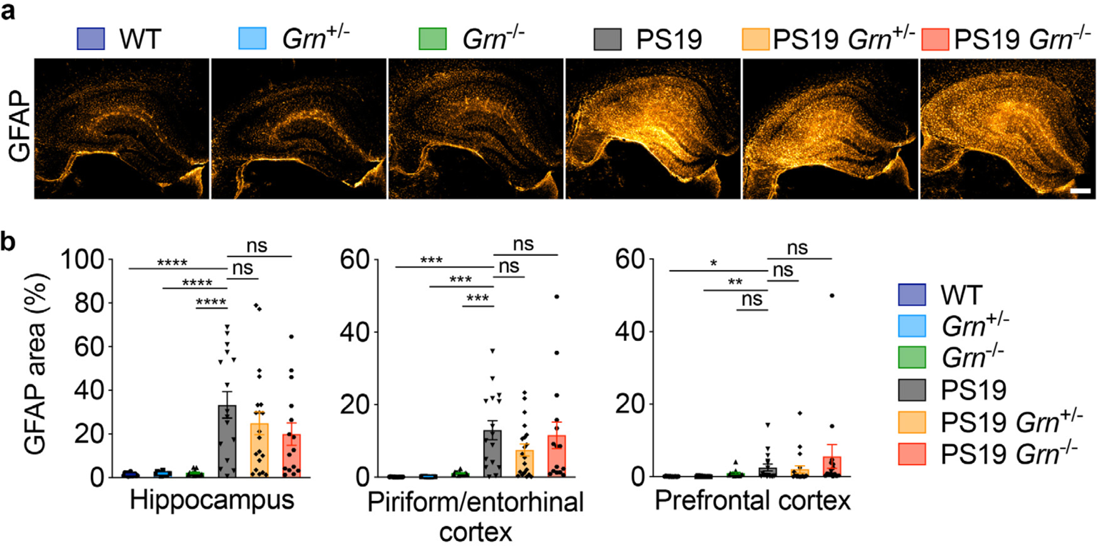
PGRN deficiency has no significant effects on astrogliosis. **a.** Representative images of GFAP staining of the hippocampus of 6 genotypes (WT, *Grn*^+/−^, *Grn*^−/−^, PS19, PS19 *Grn*^+/−^, and PS19 *Grn*^−/−^ mice) at 9-12 months of age. Bar, 300 μm. **b.** Quantification of GFAP area (%) in 9-12-month-old 6 genotypes. Mean ± SEM, n = 12–20 mice per genotype, *p < 0.05, **p < 0.01; ***p < 0.001; ****p < 0.0001; One-way ANOVA with Dunnett’s post hoc test or Kruskal-Wallis test with Dunn’s post hoc test (for the prefrontal cortex).

**Extended Data Fig. 6:**
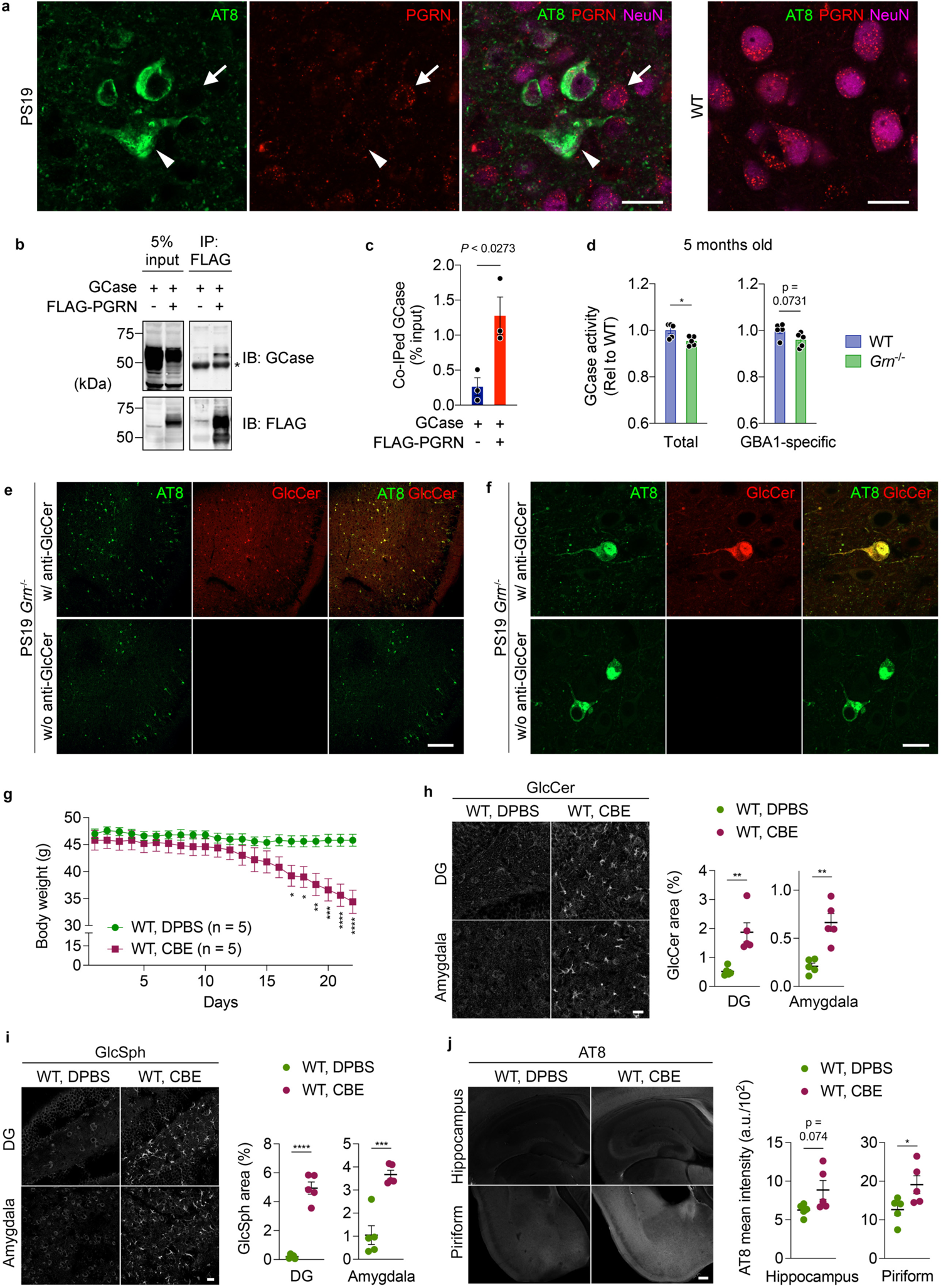
Tau inclusions are positive for GlcCer in PS19 mice with PGRN reduction. **a.** Representative confocal images of AT8, PGRN, and NeuN co-staining in the amygdala region of PS19 and WT mice. Bar, 20 μm. **b.** Co-IP assay using HEK293T cells co-expressing FLAG-PGRN and GCase suggesting interaction between PGRN and GCase. **c.** Quantification of co-IP in **a**. Mean ± SEM, n = 3 experiments, p < 0.0273; unpaired t-test. **d.** Total and GBA1-specific GCase activity of WT and *Grn*^−/−^ cortices at 5 months of age. Mean ± SEM, n = 5 mice per genotype. *p < 0.05; unpaired t-test. **e.** Representative low-magnification confocal images of AT8 and GlcCer co-staining in the piriform/entorhinal cortex region of PS19 *Grn*^−/−^ mice. In the bottom panels, the section was incubated only with AT8 antibody, followed by incubation with Alexa Fluor 488 and 568 secondary antibodies. All images were taken using the same setting. Bar, 200 μm. **f.** Representative confocal images of GlcCer and AT8 co-staining in the amygdala region of PS19 *Grn*^−/−^ mice. In the bottom panels, the section was incubated only with AT8 antibody, followed by incubation with Alexa Fluor 488 and 568 secondary antibodies. All images were taken using the same setting. Bar, 20 μm. **g.** Body weight of 8-month-old WT mice injected with DPBS or 50 mg/kg CBE every day for 21 days. Mean ± SEM, n = 5 mice per genotype. *p < 0.05, **p < 0.01, ***p < 0.001, ****p < 0.0001; two-way ANOVA with Sidak’s post hoc test. There was interaction between two factors, time and treatment (p < 0.0001). **h.** Representative images of GlcCer staining and quantification of GlcCer area (%) in the dentate gyrus (DG) and amygdala of 8-month-old WT treated with DPBS or CBE every day for 21 days. Mean ± SEM, n = 5 per groups, **p < 0.01; unpaired t-test. **i.** Representative images of GlcSph staining and quantification of GlcSph area (%) in the dentate gyrus (DG) and amygdala of 8-month-old WT treated with DPBS or CBE every day for 21 days. Mean ± SEM, n = 5 per groups, ***p < 0.001, ****p < 0.0001; unpaired t-test. **j.** Representative images of AT8 staining and quantification of AT8 mean intensity in the hippocampus and piriform cortex of 8-month-old WT treated with DPBS or CBE every day for 21 days. Mean ± SEM, n = 5 per groups, *p < 0.05; unpaired t-test.

**Extended Data Fig. 7:**
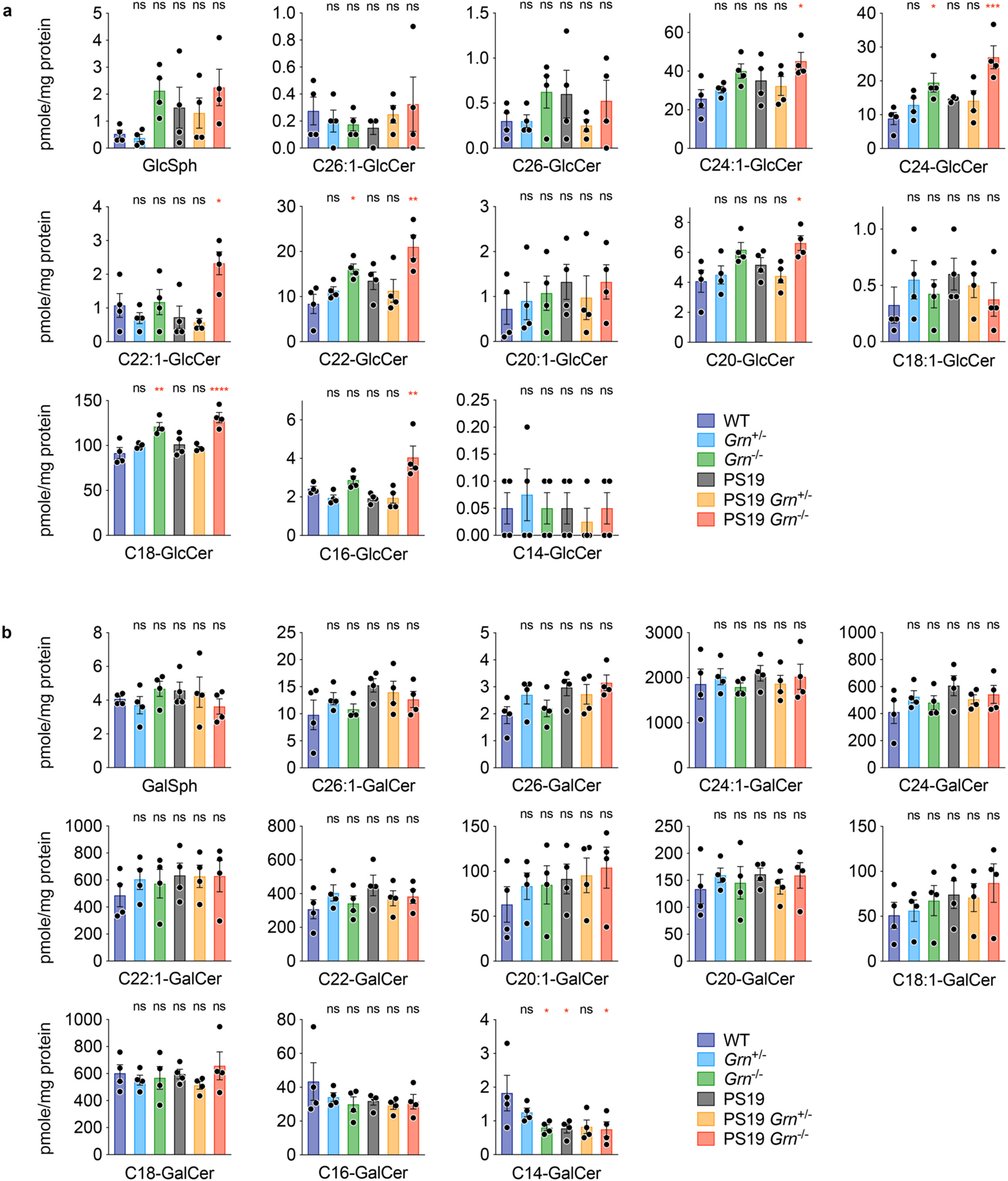
PGRN reduction increases GlcCer but not GalCer species. **a.** Levels of GlcCer species in the cortex of 6 genotypes at 10 months of age. Mean ± SEM, n = 4 mice per genotype. *p < 0.05, **p < 0.01, ***p < 0.001, ****p < 0.0001; One-way ANOVA with Dunnett’s post hoc test comparing to WT. **b.** Levels of GalCer species in the cortex of 6 genotypes at 10 months of age. Mean ± SEM, n = 4 mice per genotype. *p < 0.05; One-way ANOVA with Dunnett’s post hoc test comparing to WT.

**Extended Data Fig. 8:**
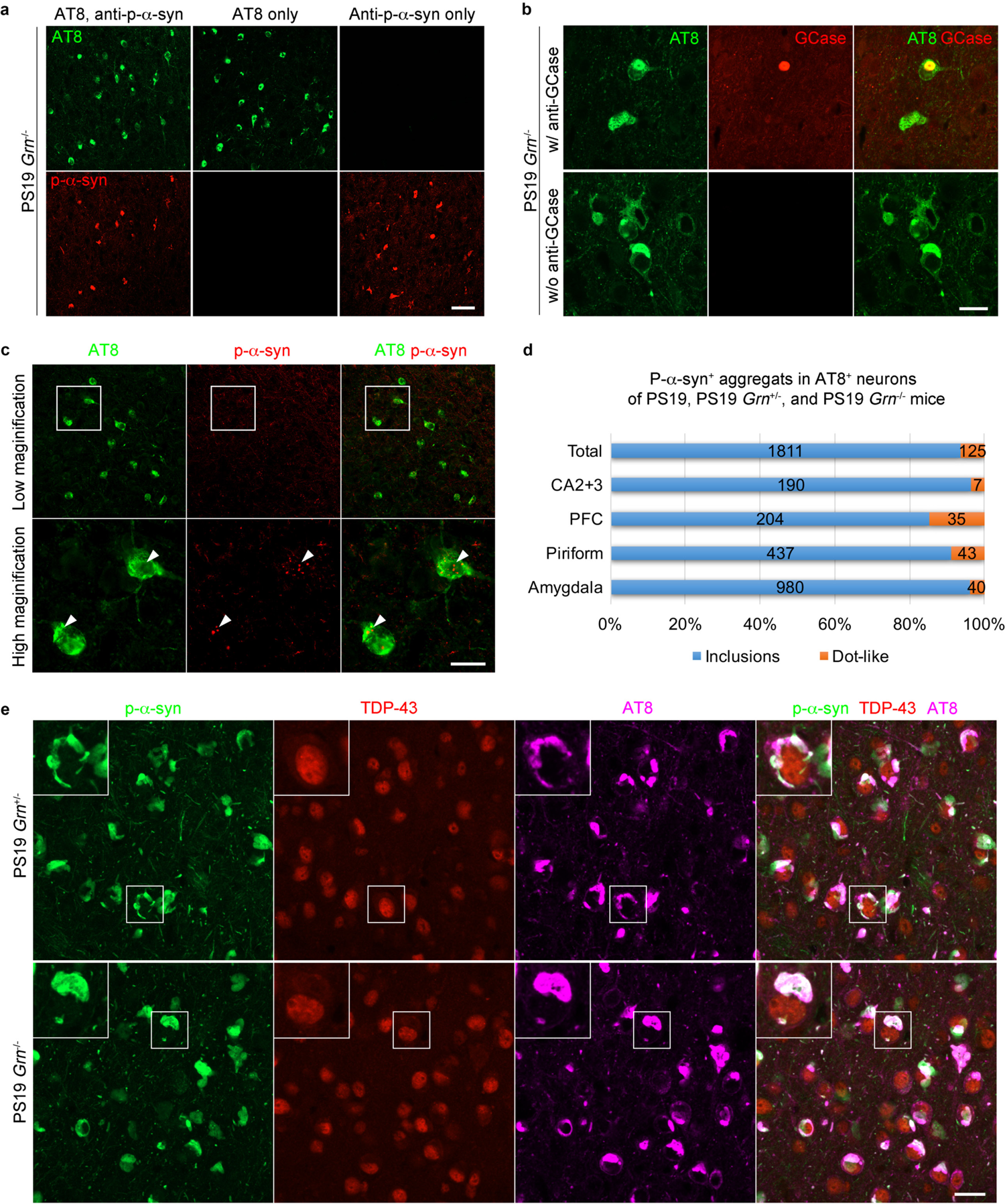
Co-localization of GlcCer and p-*α*-syn, but not TDP-43, is found in tau pathology of PS19 mice and human AD brains. **a.** Representative confocal images of AT8 and p-*α*-syn co-staining in the amygdala region of PS19 *Grn*^−/−^ mice. In the middle and right panels, the sections were incubated only with AT8 or p-*α*-syn antibody, respectively, followed by incubation with Alexa Fluor 488 and 568 secondary antibodies. All images were taken using the same setting. Bar, 50 μm **b.** Representative confocal images of AT8 and GCase co-staining of amygdala region of PS19 and PS19 *Grn*^−/−^ mice. In the bottom panels, the section was incubated only with AT8 antibody, followed by incubation with Alexa Fluor 488 and 568 secondary antibodies. All images were taken using the same setting. Bar, 20 μm. **c.** Representative confocal images of AT8 and p-*α*-syn co-staining in the amygdala of PS19 *Grn*^+/−^ mice. The top panels were imaged using 20X lens, and the bottom ones were the white rectangular area imaged using 40X lens. Arrowheads indicate dot-like aggregates of p-*α*-syn in AT8-positive neurons. Bar, 20 μm. **d.** The percentage of two types (inclusions and dot-like) of p-*α*-syn aggregates in AT8-positive neurons in each region of PS19 *Grn* cohorts. The total number of aggregates examined in each region was described in the bars. **e.** Representative confocal images of p-*α*-syn, TDP-43, and AT8 triple staining in the amygdala of PS19 *Grn*^+/−^ and PS19 *Grn*^−/−^ mice. Bar, 20 μm.

**Extended Data Fig. 9:**
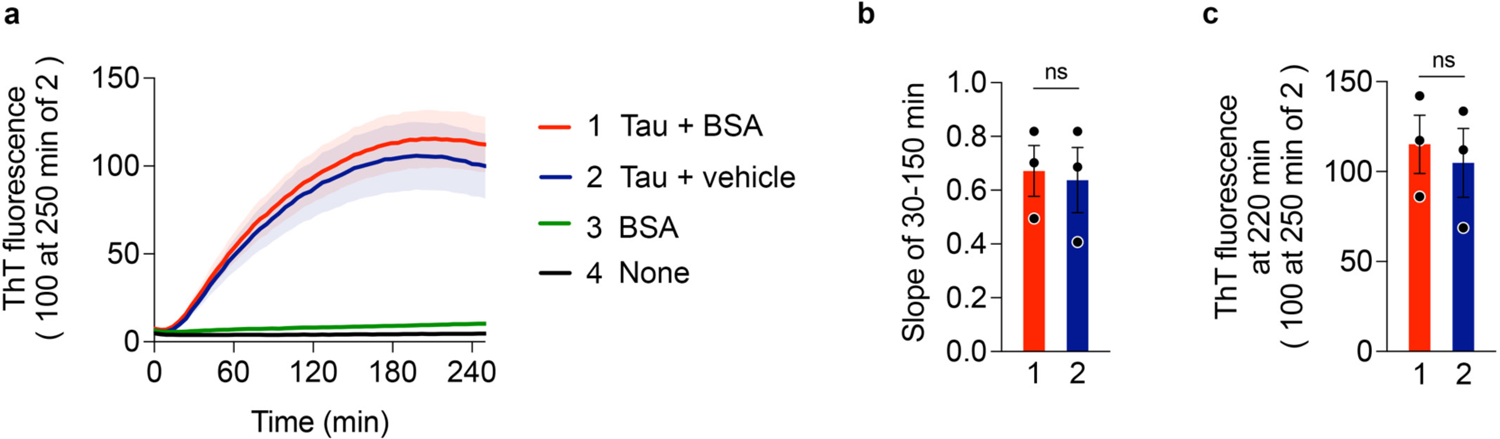
BSA has no significant effects on tau aggregation *in vitro*. **a.** ThT assay of heparin-induced aggregation of P301S tau in the presence or absence of BSA. Heparin was included in all samples except for the sample #4. Mean ± SEM, n = 3 experiments, each preformed in triplicate. **b.** Slopes determined by linear regression from 30 to 150 min in **a.** Paired t-test. **c.** ThT fluorescence at 220 min in **a.** Paired t-test.

**Extended Data Fig. 10:**
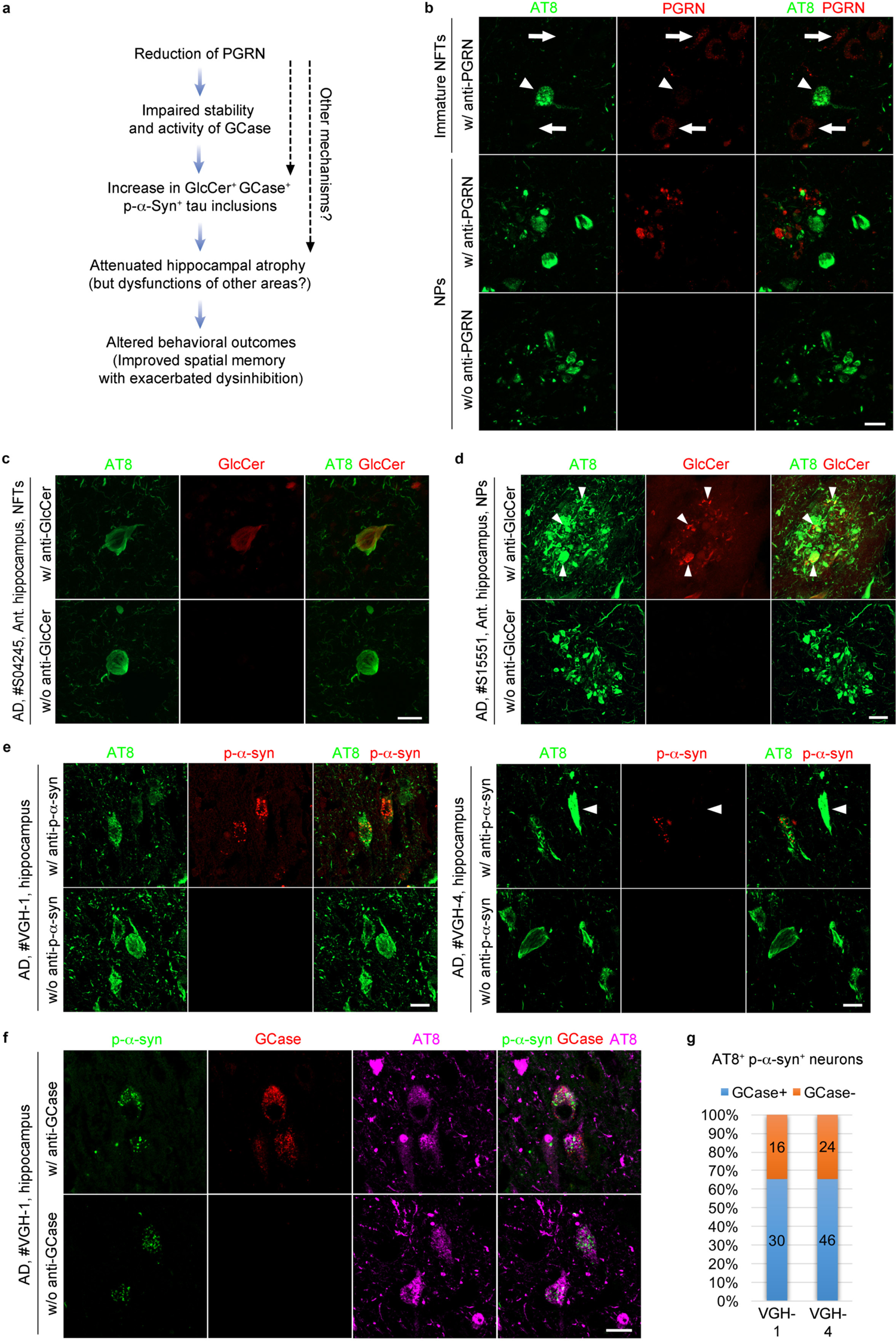
Co-localization of GCase and p-*α*-syn, but not PGRN, is found in tau pathology of human AD brains. **a.** Hypothesis of effects of PGRN reduction in tauopathy. Reduction of PGRN causes impaired activity and stability of GCase, leading to an increase in GlcCer^+^ p-*α*-syn^+^ GCase^+^ tau aggregates. Other mechanisms might be also involved in the increase of co-aggregation. The increase of co-aggregation attenuates hippocampal atrophy but might cause dysfunctions of other areas, leading to altered behavioral outcomes exhibiting exacerbated FTD-like disinhibition but improved AD-like spatial memory. It is possible that reduction of PGRN alters neurodegeneration and behaviors by other unknown mechanisms. **b.** Representative confocal images of AT8 and PGRN co-staining of immature NFTs and NPs in the hippocampus of AD brains. Arrows indicate lysosomal PGRN in neurons. An arrowhead indicates an immature NFT with very low PGRN immunoreactivity. In the bottom panels, the section was incubated only with AT8 antibody, followed by incubation with Alexa Fluor 488 and 568 secondary antibodies. All images were taken using the same setting. Bar 20 μm. **c.** Representative confocal images of AT8 and GlcCer co-staining of NFTs in the anterior hippocampus of AD patient ID#S04245. In the bottom panels, the section was incubated only with AT8 antibody, followed by incubation with Alexa Fluor 488 and 568 secondary antibodies. All images were taken using the same setting. Bar, 20 μm. **d.** Representative confocal images of AT8 and GlcCer co-staining of Nps in the anterior hippocampus of AD patients ID#S15551. In the bottom panels, the section was incubated only with AT8 antibody, followed by incubation with Alexa Fluor 488 and 568 secondary antibodies. All images were taken using the same setting. Bar, 20 μm. **e.** Representative confocal images of AT8 and p-*α*-syn co-staining of NFTs in the hippocampus of AD brain ID#VGH-1 and ID#VGH-4. An Arrowhead indicates a mature NFT with low p-*α*-syn immunoreactivity. In the bottom panels, the section was incubated only with AT8 antibody, followed by incubation with Alexa Fluor 488 and 568 secondary antibodies. All images were taken using the same setting. Bar, 20 μm. **f.** Representative confocal images of AT8, GCase, and p-*α*-syn triple staining of NFTs in the hippocampus of AD brain ID#VGH-1. In the bottom panels, the section was incubated with AT8 and anti-p-*α*-syn antibodies but without anti-GCase antibody, followed by incubation with Alexa Fluor 488, 568, and 647 secondary antibodies. All images were taken using the same setting. Bar, 20 μm. **g.** The percentage of GCase positivity in AT8^+^ p-*α*-syn^+^ neurons in AD brains #VGH-1 and #VGH-4. The number of aggregates examined is described in the bars.

**Extended Data Table 1.** Demographic information for human samples used in this study

| Brain ID | Diagnosis | Age | Sex | PMI | Braak | Region examined | Section | Source |
| --- | --- | --- | --- | --- | --- | --- | --- | --- |
| S05630 | AD | 78 | F | 2.58 | VI | Anterior hippocampus | Free-floating | HBTRC |
| S15551 | AD | 82 | F | 16.75 | V | Anterior hippocampus | Free-floating | HBTRC |
| S15147 | AD | 87 | F | 6.17 | V | Anterior hippocampus | Free-floating | HBTRC |
| S04245 | AD | 63 | F | 9 | V-VI | Anterior hippocampus | Free-floating | HBTRC |
| S19475 | AD | 82 | M | 24.8 | V-VI | Anterior hippocampus | Free-floating | HBTRC |
| S08380 | AH2 | 55 | M | 33 | III | Anterior hippocampus | Free-floating | HBTRC |
| S18596 | AH2 | 100 | F | 3.16 | III | Anterior hippocampus | Free-floating | HBTRC |
| S15386 | AH2 | 89 | M | 14 | III | Anterior hippocampus | Free-floating | HBTRC |
| S06423 | AH2 | 84 | M | 7.31 | III-IV | Anterior hippocampus | Free-floating | HBTRC |
| S17240 | DLB | 67 | M | 14.85 | 0 | Anterior hippocampus | Free-floating | HBTRC |
| S19496 | DLB | 80 | M | 16.33 | II | Anterior hippocampus | Free-floating | HBTRC |
| S13303 | DLB | 68 | M | 5.5 | I-II | Anterior hippocampus | Free-floating | HBTRC |
| S16963 | DLB | 80 | M | 25.25 | II | Anterior hippocampus | Free-floating | HBTRC |
| 13-78 | FTD-GRN | 72 | F | 6.45 | VI | Middle frontal gyrus | Free-floating | BSHRI |
| VGH-1 | AD | 73 | F | 48 | VI | Hippocampus | Paraffin-embedded | VGH |
| VGH-2 | AD | 71 | M | 24 | VI | Hippocampus | Paraffin-embedded | VGH |
| VGH-3 | AD | 90 | F | 48 | VI | Hippocampus | Paraffin-embedded | VGH |
| VGH-4 | AD | 83 | F | 48 | VI | Hippocampus | Paraffin-embedded | VGH |
| VGH-5 | FTD-GRN | 77 | F | 12 | III | Hippocampus | Paraffin-embedded | VGH |
| VGH-6 | FTD-GRN | 85 | F | 12 | II | Hippocampus | Paraffin-embedded | VGH |

AH2; Alzheimer's changes moderate
HBTRC; Harvard brain tissue resource center
BSHRI; Banner Sun Health Research Institute
VGH; Vancouver General Hospital

### Online Methods

#### Lead contact

Further information and requests for resources and reagents should be directed to and will be fulfilled by the Lead Contact, Stephen M. Strittmatter.

#### Data and code availability

snRNA-seq raw data have been deposited at GEO and are publicly available as of the date of publication. Accession numbers are listed in the key resources table. This paper does not report new unique codes. Any additional information required to reanalyze the data reported in this paper is available from the lead contact upon request.

#### Animals

Human P301S tau transgenic (PS19) mice with B6C3F1 background ^34^ were obtained from the Jackson Laboratory. The mice express mutant human *MAPT* gene, which results in a five-fold greater amount of mutant human tau protein than endogenous mouse tau. *Grn*^−/−^ mice with C57BL/6J background ^35^ were obtained from the RIKEN Bioresource Center. To generate 6 genotypes used in this study (WT, *Grn*^+/−^, *Grn*^−/−^, PS19, PS19 *Grn*^+/−^, and PS19 *Grn*^−/−^) with minimal differences in genetic background, PS19 and *Grn*^−/−^ mice were first crossed to generate PS19 *Grn*^+/−^ and *Grn*^+/−^ mice and then these mice were crossed. For behavioral tests, both male and female mice were used with similar ratio between genotypes at 10-11 months of age. For volumetric, immunohistochemcal, and biochemical analyses, only male were used at 9-12 months of age (mean ages are not significantly different between genotypes in all analyses) as previous studies have reported significantly more tau pathology and neurodegeneration in male versus female PS19 mice ^42–45^.

Animals were housed in groups with 2-5 animals per cage and maintained on 12 h light-dark schedule with access to food and water *ad libitum.* All protocols were approved by Yale Institutional Animal Care and Use Committee (IACUC).

#### Conduritol B Epoxide (CBE) treatment in mice

GCase inhibitor CBE (MedChemExpress #HY-100944) was reconstituted at 10 mg/mL in DPBS and stored at –80°C. Eight-months-old male WT mice with B6C3F1 background were injected intraperitoneally with 50 mg CBE per kg body weight or vehicle (DPBS) per day, for 21 days, alternating sides of injection.

#### Human brain tissue

Samples of pre-existing human autopsy brain with no personal identifiers were obtained from Harvard Brain Tissue Resource Center, Banner Sun Health Research Institute, and Vancouver General Hospital (Table S1).

#### Primary neuronal culture

Primary mouse hippocampal and cortical neurons were prepared from E16-18 embryos (both male and female) as described previously ^67^, plated at 75,000 cells/well onto poly-D-lysine (PDL)-coated 96 well plates (CORNING #354461) and cultured in Neurobasal-A medium (GIBCO #10888-022) supplemented with B27 supplement (GIBCO #17504-044), 1 mM sodium pyruvate (GIBCO #11360-070), GlutaMAX-I (GIBCO #35050-061), and 100 U/mL penicillin and 100 mg/mL streptomycin (GIBCO #15140-122) at 37°C in 5% CO_2_.

#### Cell culture and transfection

HEK293T cells (ATCC #CRL-11268) were maintained in DMEM (GIBCO #11965-092) supplemented with 10% fetal bovine serum (GIBCO #16000-044), 100 U/mL penicillin and 100 mg/mL streptomycin (GIBCO #15140-122) at 37°C in 5% CO_2_. Transient transfection was performed with Lipofectamine 2000 (Invitrogen #11668-019) according to the manufacturer’s protocol.

#### Mouse behavioral tests

All behavioral tests were conducted in a blinded manner. Both male and female were used at 10-11 months of age. Prior to behavioral tests, each mouse was handled for 5 min for 4 days to reduce anxiety. In all behavioral tests, mouse behavior was recorded on a JVC Everio G-series camcorder (Yokohama, Japan) and tracked by Panlab SMART software (Barcelona, Spain).

#### Open field test

Open field test was performed as previously described ^39^ with slight modifications. Briefly, the apparatus for the test consists of a gray, 50 cm wide, 50 cm long, 40 cm high acrylamide box, with an open top. Two light sources were put by east and west sides of the box. Individual mice were placed at the corner of the box and monitored for 10 min. The bottom of box was cleaned with 70% ethanol after every trial.

#### Elevated plus maze

Elevated plus maze (EPM) was performed as previously described ^39^. Briefly, EPM was set at a height of 65 cm and consisted of two open white Plexiglas arms, each arm 8 cm wide x 30 cm long and two enclosed arms (5 cm x 30 cm) with black 15 cm high walls that were connected by a central platform (5 cm x 5 cm). Individual mice were placed at the central platform and monitored for 5 min. All arms were cleaned with 70% ethanol after every trial.

#### Morris water maze

Morris water maze (MWM) was performed as previously described ^26, 67^ with slight modifications. MWM was performed at room temperature and water temperature was kept at 20-25°C. Mice were placed in an ∼ 1 meter diameter pool with a hidden, clear platform filled with water to 1 inch above the submerged platform. The hidden platform was placed in one of the 4 quadrants of the pool with the 4 drop zones directly across from the platform. A symbol, such as a plus or a cross, was placed at each of the 4 cardinal directions as possible recognition flags. The learning trial consisting of 6 trail blocks was performed two times per day for 3 consecutive days. Mice were dropped off facing the wall at 4 different drop zones (four trials in each trial block). Each trial block was performed by alternating two mice (e.g. A, B, A, B, A, B, A, B, C, D, C, D…). Latency was measured as the time that it took for the mouse to find and spend 1 second on the hidden platform. If there was a failure to reach the platform in 60 s, the mouse was guided to the platform and allowed to rest on it for 15 s. The probe trial was performed ∼24 h after finishing the learning trail. During the probe trail, the platform was removed from the pool and mice were started from a location along the pool wall diagonally opposed to the location of the platform in the learning trail and allowed to swim in the pool for 60 s. Latencies for the learning and probe trials were measured with the Panlab SMART Video Tracking Software (ver 2.5.21).

After the probe trial, the visible platform trial was performed, placing a flag (50 mL tube) atop of the hidden platform. The trial was performed by alternating two mice and mice were repeatedly placed in the pool. Time taken to reach the visible platform was manually recorded. When a consistent time for a mouse was reached, the last three times were averaged.

#### Brain tissue collection and processing

Mice were euthanized with CO_2_ and perfused with ice-cold PBS for 80 s. The brains were dissected out and the hemispheres were divided. The left hemisphere was immediately snap frozen in liquid nitrogen or the hippocampus and cortex were dissected from the left hemisphere and were individually snap frozen for biochemical analysis. The right hemisphere was fixed in 4% paraformaldehyde (PFA) (SIGMA #158127) in PBS for 1 day at 4°C and then embedded in 10% gelatin (SIGMA #G1896) and placed in 4% PFA for another 3 days at 4°C. Fifty µm coronal sections of the right hemisphere were cut with a Leica VT1000S Vibratome and stored in PBS with 0.05% sodium azide at 4°C for volumetric and immunohistochemical analyses.

#### Volumetric analysis and neuronal layer thickness measurement

Volumetric analysis and neuronal layer thickness measurement were performed as previously described ^43^ with modifications. Every sixth section (0.3 mm between sections) from bregma −1.1 mm to −3.9 mm was mounted for each mouse. After drying overnight, all sections were stained with 0.1% cresyl violet (SIGMA #C5042) for 10 min at 50°C. The slides were then rinsed with H_2_O for 1 min, sequentially dehydrated in 95% ethanol for 15 min (twice) and 100% ethanol for 5 min (twice), cleared in xylene for 5 min (twice), and coverslipped in CYTOSEAL60 (ThermoFisher #8310-4). The stained sections were imaged using Aperio CS2 scanner (Leica) at 20X magnification and analyzed using ImageScope software. For hippocampus and posterior lateral ventricle, sections between bregma −1.1 mm and −3.9 mm were used for quantification. For piriform/entorhinal cortex, sections between bregma −2.3 mm and −3.9 mm were used for quantification. The volume was calculated using the following formula: (sum of area) x 0.3 mm.

Neuronal layer thickness measurement was performed using three sections (bregma −1.4, −1.7, and −2.0 mm) that were stained with cresyl violet as described above. The thickness was measured using the Distance Measurement tool in ImageScope software. All staining and data analysis were performed by a researcher who was blinded to the genotype.

#### Immunohistochemistry

Immunohistochemistry was performed as previously described ^26^ with slight modifications. For AT8/GFAP co-staining, five sections (approximately bregma 2.0, and 1.7 mm for prefrontal cortex, −1.4, −1.7, and −2.0 mm for hippocampus and piriform/entorhinal cortex) were used. For Iba1/CD68 co-staining, four sections (approximately bregma 2.0 and 1.7 mm for prefrontal cortex, −1.4 and −1.7 mm for hippocampus and piriform/entorhinal cortex) were used. For PGRN/Iba1 co-staining, two sections (approximately bregma −1.4 and −1.7 mm) were used. For AT8/p-*α*-syn co-staining, two sections (approximately bregma 1.9 and −2.4 mm) were used. Free-floating sections were permeabilized and blocked with 10% normal donkey serum, 0.2% Triton X-100 in PBS for 1 h at room temperature. The sections were then incubated in primary antibody in 1% normal donkey serum, 0.2% Triton X-100 in PBS overnight at room temperature. The primary antibodies that were used include: mouse PGRN (R&D #AF2557, 1:400), human PGRN (R&D #AF2420, 1:200), AT8 (Invitrogen #MN1020, 1:500), CD68 (Bio-Rad #MCA1957, 1:900), Iba1 (FUJIFILM Wako #019-19741, 1:500), GFAP (Abcam #ab7260, 1:1000), GlcCer (Glycobiotech #RAS_0011, 1:100), GCase (SIGMA #G4171, 1:150), phospho-*α*-synuclein (BioLegend #MMS-5091, 1:250), TDP-43 (proteintech #10782-2-AP, 1:250). The sections were then washed three times with PBS and then incubated for 2-3 h at room temperature in either donkey anti-rabbit or donkey anti-mouse fluorescent secondary antibodies (Invitrogen Alexa Fluor 1:500) in 1% normal donkey serum, 0.2% Triton X-100 in PBS. For AT8/p-*α*-syn co-staining, normal goat serum was used for blocking and dilution of antibodies, and goat anti-mouse IgG2a, Alexa Fluor 568 (Invitrogen #A-21134) and goat anti-mouse IgG1, Alexa Fluor 488 (Invitrogen #A-21121) secondary antibodies were used. For AT8/p-*α*-syn/TDP-43 triple staining, normal goat serum was used for blocking and dilution of antibodies, and goat anti-mouse IgG2a, Alexa Fluor 488 (Invitrogen #A-21131), goat anti-rabbit IgG, Alexa Fluor 568 (Invitrogen #A11036), and goat anti-mouse IgG1, Alexa Fluor 647 (Invitrogen #A-21240) secondary antibodies were used. After incubation, the sections were washed three times with PBS. To quench autofluorescence, sections were dipped briefly in dH_2_O and then incubated in 10 mM copper sulfate, 50 mM ammonium acetate, pH 5 for 15 min before dipping back into dH_2_O and then placed in PBS ^80^. All sections were mounted onto glass slides (Superfrost plus, Fischer Scientific Company L.L.C.) and coverslipped with Vectashield antifade mounting medium with DAPI (Vector). All procedures were performed by a researcher who was blinded to the genotype.

#### Immunohistochemistry with human brain samples

Formalin-fixed brains from Harvard and Banner were cut with Leica VT1000S Vibratome, and free-floating sections (50 μm) were stored in PBS with 0.05% sodium azide at 4°C. For the free-floating sections, antigen retrieval was performed by autoclaving at 120°C for 10 min in 10 mM sodium citrate buffer, pH 6.0 before performing the immunohistochemical procedure described above. Paraffin-embedded sections from Vancouver were deparaffinized, rehydrated, and incubated in preheated 10 mM sodium citrate buffer, pH 6.0 for 30 min at 95°C before performing the immunohistochemical procedure described above. SUPER PAP PEN (EMS #71312) was used to perform the procedure with minimum volume of antibody solution. Humidity was kept by using a storage container (Rosti Mepal, 500 mL) with wet paper towel on the bottom during the procedure. For AT8/p-*α*-syn/GCase triple staining, normal goat serum was used for blocking and dilution of antibodies, and goat anti-mouse IgG2a, Alexa Fluor 488 (Invitrogen #A-21131), goat anti-rabbit IgG, Alexa Fluor 568 (Invitrogen #A11036), and goat anti-mouse IgG1, Alexa Fluor 647 (Invitrogen #A-21240) secondary antibodies were used. To check background and bleed-through signals, a control staining without the primary antibody was performed side by side for each sample and antigen.

#### Imaging

For AT8/GFAP, Iba1/CD68, and PGRN/Iba1 staining, all images of hippocampus, piriform/entorhinal cortex, and prefrontal cotex were taken using a Zeiss AxioImager Z1 fluorescent microscopy with a 5X objective lens. For AT8/PGRN, AT8/GlcCer, AT8/p-*α*-syn, and AT8 or p-*α*-syn/GCase staining, images were taken using Zeiss LSM800 confocal microscopy with 10X, 20X, 40X or 63X objective lens. For AT8/p-*α*-syn co-staining, three confocal images with higher number of AT8 and p-*α*-syn double positive inclusions were taken from each region (prefrontal cortex, CA2+3, amygdala, piriform cortex) with 20X objective lens. For p-*α*-syn/GCase images in Figure 5 the maximal intensity projection function was used with z-stack confocal images.

#### Nuclei isolation from the mouse hippocampus

Nuclei were isolated with Nuclei EZ Prep Kit (SIGMA #NUC-101) as previously reported ^81^ with modifications. Frozen hippocampus was homogenized using a glass dounce tissue grinder (SIGMA #D8938) 25 times with pastel A and 25 times with pastel B in 2 mL of ice-cold lysis buffer (Nuclei EZ lysis buffer with Complete Mini EDTA-free and Protector RNase Inhibitor (ROCHE)) and transferred to 15 mL tube with additional 2 mL of ice-cold lysis buffer, and then vortexed and incubated on ice for 5 min. Nuclei were centrifuged at 500 x g for 5 min at 4°C, washed with 4 mL of ice-cold lysis buffer, briefly vortexed and incubated on ice for 5 min. After centrifugation at 500 x g for 5 min at 4°C, the nuclei were washed with 4 mL of ice-cold nuclei suspension buffer (0.5 mg/mL BSA (ThermoFisher #AM2616) and 0.1% Protector RNase inhibitor in DPBS (CORNING #21-031-CV)) once, resuspended in 1 mL of nuclei suspension buffer, filtered through a 35 μm cell strainer (FALCON #352235) twice, and counted with trypan blue. The samples were diluted to a final concentration of ∼1,000 nuclei per μL and used for subsequent analyses. Nuclei of three samples ((WT, *Grn*^+/−^, and *Grn*^−/−^) or (PS19, PS19 *Grn*^+/−^, and PS19 *Grn*^−/−^)) were isolated and submitted for the sequencing at a time and the process was repeated six times to obtain three biological replicates for each genotype.

#### Single-nucleus RNA sequencing (snRNA-seq)

Libraries were prepared using Chromium Single Cell 3’ GEM, Library & Gel Bead Kits v3 (10X Genomics #PN-1000075) according to the manufacturer’s protocol. The libraries underwent sequencing with a sequence depth of 200 million reads per sample using NovaSeq instrument (Ilumina) according to 10X Genomics recommendations. Sample demultiplexing, alignment, and gene counting were performed using Cell Ranger software (10X Genomics, version 3.0.2). Gene counts were obtained by aligning reads to the mm10 reference genome.

#### Quality control and pre-processing of snRNA-seq datasets

Subsequent data processing was performed using the Seurat package (version 3.1.4) ^82^. In all 18 datasets, genes expressed in < 3 nuclei and nuclei with > 5% mitochondrial counts were removed. Nuclei that express < 200 or > 2,500 genes were also removed for downstream analysis.

Doublets were detected and removed using the DoubletFinder package (version 2.0.3) prior to dataset integration ^83^. All datasets were pre-processed with SCTransform ^84^, RunPCA (principal component analysis), and RunUMAP (Uniform Manifold Approximation and Projection) functions using the Seurat package. The proportion of artificial doublets (pN) was set to 0.25 (authors’ recommendation) in all datasets, and the neighborhood size parameter (pK) was optimized for each dataset with the mean-variance-normalized bimodality coefficient (BCMVN) maximization as described previously ^83^. The doublet rate was set to 0.075 based on 10x Genomics protocol without homotypic doublet adjustment to remove all potential heterotypic doublets.

#### Integration, visualization, and clustering of snRNA-seq datasets

In order to reduce potential batch effects, “Reference-based” integration was performed with 18 SCTransform-normalized and doublet-removed datasets using the Seurat package as shown in online vignettes (https://satijalab.org/seurat/v3.1/integration.html).

A reference was selected from each genotype and total 6 references were used for the integration. For visualization and clustering, the integrated dataset was processed with RunPCA and RunUMAP (with the top 50 PCs) functions, followed by FindNeighbors and FindClusters (with a resolution of 0.3) functions of the Seurat package. These analyses identified 33 pre-clusters in the integrated dataset. Cell types were then assessed for each pre-cluster based on the expression of known maker genes and previously identified signatures using the hippocampal samples ^49, 81, 85^. Pre-clusters that were defined to be the same cell type were grouped and a pre-cluster with nuclei with low UMI counts was removed for downstream analysis. Due to the low number of nuclei, cell type could not be defined in one pre-cluster (cluster #13). The resulting 12 cell type clusters were used for differential expression and enrichment analyses.

#### Brain extraction

Sequential fractionation using RAB, RIPA, 70% formic acid (FA) buffer was performed as described previously^43^ with modifications. Briefly, cortices from left hemisphere were weighted and homogenized using a dounce homogenizer (DWK Life Science #357422) for 25 strokes in 10-fold volume of ice-cold RAB buffer (100 mM MES, 1 mM EGTA, 0.5 mM MgSO_4_, 750 mM NaCl, 20 mM NaF, 1 mM Na_3_VO_4_, pH 7.0, supplemented with PhosSTOP, cOmplete-mini (Roche)). After ultracentrifugation for 20 min at 50,000 x g at 4°C, the supernatant was collected and saved as the RAB-soluble fraction and the pellet was resuspended in 10-fold volume of ice-cold RIRA buffer (25 mM Tris, 150 mM NaCl, 1% NP40, 0.5% deoxycholic acid, 0.1% SDS, 20 mM NaF, 1 mM Na_3_VO_4_, pH 8.0, supplemented with PhosSTOP, cOmplete-mini (Roche)) and nutated for 30 min at 4°C. After ultracentrifugation for 20 min at 50,000 x g at 4°C, the supernatant was collected and saved as the RIPA-soluble fraction and the pellet was resuspended in 3-fold volume of ice-cold 70% FA and nutated for 30 min at 4°C. After ultracentrifugation for 20 min at 50,000 x g at 4°C, the supernatant was collected and saved as the FA-soluble fraction. All fractions were stored at –80°C.

#### Co-immunoprecipitation assay

Co-immunoprecipitation (co-IP) assay was performed as previously reported ^39^ with modifications. Plasmids encoding non-tagged human GBA were purchased from OriGene Technologies, Inc. FLAG-PGRN was described previously ^86^. One day after transfection, HEK293T cells were harvested and lysed with ice-cold 50 mM Tris-HCl pH 7.5, 150 mM NaCl, 0.5% Triton X-100 supplemented with PhosSTOP, cOmplete-mini (Roche). After centrifugation for 30 min at 100,000 x g at 4°C, the supernatant was incubated with anti-FLAG M2 Affinity Gel (SIGMA #A2220) overnight at 4°C. The immunoprecipitates were washed five times with ice-cold 50 mM Tris-HCl pH 7.5, 150 mM NaCl, 0.1% Triton X-100 and boiled with 2 x Laemmli sample buffer (Bio-Rad) with *β*ME.

#### Immunoblot

Immunoblot was performed as previously reported ^26^, with slight modifications. For the FA-soluble fraction, the samples were diluted with 16-fold volume of 1 M Tris. The protein samples were electrophoresed using precast 4-20% Tris-glycine gels (Bio-Rad) and transferred with an iBlot 2 Transfer Device onto nitrocellulose membranes (Invitrogen IB23001). The membranes were incubated in blocking buffer (Rockland MB-070) for 1 h at room temperature and then incubated overnight at 4°C in blocking buffer with primary antibodies: GCase (SIGMA #G4171, 1:1000) and FLAG (SIGMA #F7425, 1:1000), tau (DAKO #A0024, 1:5000), HT7 (Invitrogen #MN1000, 1:1000), AT8 (Invitrogen #MN1020, 1:1000), PHF1 (gift from Dr. Peter Davies, Albert Einstein College of Medicine, Bronx, NY, 1:1000), phospho-tau (Ser199, Ser202) (Invitrogen #44-768G, 1:1000), phospho-tau (Ser356) (Invitrogen #44-751G, 1:1000), *β*-actin (Cell Signaling Technology #3700, 1:2000). The next day, the membranes were washed three times with TBST for 3 min and incubated in secondary antibodies (Li-Cor, IR Dye 680 or 800, all 1:10,000) for 1 h at room temperature. After washing three times with TBST for 3 min, proteins were visualized with an Odyssey Infrared imaging system (Li-Cor). The immunoreactive bands were quantified using ImageJ software.

#### GCase activity assay

GCase activity assay was performed as previously reported ^55, 56^ with modifications. Cortices from the left hemispheres were homogenized in 10-fold volume of ice-cold citrate-phosphate buffer pH 5.2, 1% Triton X-100 with PhosSTOP and cOmplete-mini (Roche) using a dounce homogenizer (DWK Life Science #357422) for 20 strokes and nutated for 30 min at 4°C. After ultracentrifugation for 30 min at 100,000 x g at 4°C, the supernatants were collected and stored at –80°C. On the day of the experiment, the supernatants were thawed and 5 μL of each sample was pre-incubated in a black 96-well plate (Costar #3916) for 30 min on ice with 10 μL of 2.5 mM Conduritol B Epoxide (CBE) in GCase activity assay buffer (citrate-phosphate buffer pH 5.2, 1% BSA (SIGMA #A9647), 4 mM sodium taurocholate (SIGMA #T4009), 150 μM EDTA) and then mixed with 10 μL of 10 mM 4-Methylumbelliferyl *β*-D-glucopyranoside (SIGMA #M3633) in GCase activity assay buffer for a total reaction volume of 25 μL. All assays were performed in duplicate. Reactions were performed on ice for 30 min with brief shaking every 5 min and then at 37°C for 60 min. Reactions were stopped by adding 200 μL of precooled 0.5 M glycine-NaOH, pH 10.6 stop solution on ice. Measurements were taken with the Victor 3 plate reader (PerkinElmer) at excitation of 355 nm and emission of 460 nm. Protein concentrations were determined using Protein Assay Dye Reagent Concentrate (BIO-RAD #5000006) and used for normalization.

#### Lipidomics

Separation of GlcCer and GalCer species was performed at the Lipidomics Shared Resource at Medical University of South Carolina. Cortices were weighed and homogenized in 10-fold volume of ice-cold Tissue Homogenization Buffer (0.25 M sucrose, 25 mM KCl, 0.5 mM EDTA, 50 mM Tris-HCl, pH 7.4). Lipids were extracted from homogenate containing 1 mg of protein, and levels of GlcCer and GalCer species were measured with supercritical fluid chromatography-tandem mass spectrometry analysis.

### AD-tau seeding assay

AD-tau seeding assay was performed as previously reported ^67, 68^, with slight modifications. Primary cultured neurons were prepared from mouse E16-18 WT (C57BL6) or *Grn*^-/-^ embros (both male and female). Dissociated neurons from the hippocampus and cortex were plated at 75,000 cells/well onto poly-D-lysine-coated 96-well plates (Corning #354461). Experiments were performed in triplicate. At DIV7, tau fibrils extracted from AD patients (AD-tau) were seeded onto neurons. Conduritol B Epoxide (CBE) or vehicle (ultrapure water) was also treated with AD-tau at DIV7. At DIV21, the neurons were fixed with ice-cold methanol for 30 min on ice, and blocked with 10% normal donkey serum, 0.2% Triton X-100 in PBS for 1 h at room temperature. Neurons were then incubated overnight at 4°C in 1% normal donkey serum, 0.2% Triton X-100 in PBS with primary antibodies: Mouse tau (T49) (SIGMA, #MABN827, 1:500) and MAP2 (Cell signaling, #4542, 1:150). The next day, neurons were washed with PBS two times and incubated for 1 h at room temperature with secondary antibodies anti-mouse IgG Alexa Fluor 488 (Invitrogen, 1:500) and anti-rabbit IgG Alexa Fluor 568 (Invitrogen, 1:500) and DAPI (0.5 μg/mL) diluted in 1% normal donkey serum, 0.2% Triton X-100 in PBS. Finally, neurons were washed four times with PBS and imaged automatically and unbiasedly (4 images/well) using ImageXpress Micro XLS with 20X objective lens (Molecular Devices).

### Preparation of lipid dispersions

Lipid dispersions were prepared as previously reported ^63^ with modifications. Brain phosphatidylchorine (PC) (Avanti Polar Lipids #840053P), Glucosyl (*β*) Ceramide (GlcCer) (Avanti Polar Lipids #860547P), Glucosyl (*β*) Sphingosine (Avanti Polar Lipids #860535P), and Galactosyl (*β*) Ceramide (Avanti Polar Lipids #860844P) were dissolved in chloroform at 10 mg/mL. PC was aliquoted and stored at −20°C in glass vials with nitrogen gas overlay. The other lipids was aliquoted and used immediately or stored at −20°C in glass vials with nitrogen gas overlay and used the next day. For lipid dispersions, lipids were mixed at 1:3 molar ratio and vortexed thoroughly in glass tubes before drying under a nitrogen stream. The lipid film was hydrated in fibrillization buffer (10 mM Hepes, 100 mM NaCl, 1 mM DTT, pH 7.4), vortexed thoroughly, and sonicated in a water bath for 15 min. After transferring to polypropylene tube, the samples were bath sonicated for 15 min, put through 4 freeze/thaw cycles, and bath sonicated for another 30 min. The lipid dispersions were vortexed and then used immediately.

### Thioflavin T assay

A final concentration of 6.6 μM recombinant human P301S tau 2N4R (StressMarq #SPR-327) was mixed with 4.5 μM heparin (SIGMA #H3393) and 25 μM Thioflavin T (ThT; SIGMA #T3516) in fibrillization buffer (10 mM Hepes, 100 mM NaCl, 1 mM DTT, pH 7.4), and then with 1.2 μM recombinant human GCase (R&D system #7410-GHB), which was dissolved in 50 mM sodium citrate pH 5.5, BSA, or lipid dispersions described above (787 μM monomer equivalent). Forty μL volumes were added to black 384-well plates (Greiner 781900). All experiments were performed in triplicate. The plate was incubated for 255 min at 37°C in the Victor 3 plate reader (PerkinElmer). ThT fluorescence (at excitation of 440/8 nm and emission of 486/10 nm) was measured every 5 min with 15 s shaking in a double-orbital fashion before the measurement.

### Transmission EM

Three micro liters of diluted samples after ThT assay were placed on carbon-coated TEM grids (Electron Microscopy Sciences CF300-CU) that were glow discharged for 45 seconds. Excess material was blotted with filter paper and the grid was washed and stained in 2% uranyl acetate for 1 minute before blotting and drying. The grids were imaged either in an FEI 120 kV Tecnai TEM with an Ultrascan 4000 CCD Camera or an FEI 200 kV TF20 with an AMT Nanosprint1200 CMOS Camera. At least two independent image sessions were conducted for each sample to confirm the reproducibility.

### Image quantification of immunohistochemistry

All quantitative analyses of images taken with Zeiss AxioImager Z1 fluorescent microscopy were done using ImageJ (National Institutes of Health). For Iba1/CD68 co-staining, after background subtraction (Rolling ball radius: 200 pixels), all images were uniformly thresholded and binarized. Iba1 and CD68 areas were calculated using the “analyze particles” of ImageJ. The mean of two or three sections was used to represent each mouse. For AT8/p-*α*-syn co-staining, three images taken from each region (prefrontal cortex, CA2+3, amygdala, piriform cortex) were used to manually count AT8 and p-*α*-syn positive inclusions. All imaging and data analyses were performed by a researcher who was blinded to the genotype.

### Image quantification of AD-tau seeding assay

All quantitative analyses were performed as previously reported ^67^ using macro of ImageJ (National Institutes of Health). For the graphs with data points of “Experiments”, the mean of 12 images (3 wells x 4 images) from each sample was used to represent each experiment. Data analyses were performed by a researcher who was blinded to the group or genotype.

### Data analysis of GCase activity assay

After subtraction of background obtained from wells with GCase activity buffer containing only 4 mM 4-Methylumbelliferyl *β*-D-glucopyranoside, the mean values of duplicate wells were used to represent each sample. The values from samples without CBE were used as total GCase activity and normalized to the mean value of WT or PS19 samples. For GBA1-specific activity, values from samples with CBE were subtracted from those from sample without CBE and then the subtracted values were normalized to the mean values of WT or PS19 samples.

### Statistical analysis for behavioral tests, volumetric analysis, immunohistochemistry, and GCase activity assay

One-way ANOVA, two-way ANOVA, or Kruskal-Wallis test with multiple comparisons tests (for > 3 groups) and two-tailed unpaired t-test (for 2 groups) were performed as specified in the figure legends using GraphPad Prism 8. Assumption of whether or not data follow normal distribution was based on previous studies ^26, 39, 43^. In EPM test, outliers detected by ROUT (Q = 1%) were excluded. In MWM, outliers that took more than 30 s to reach the visible platform were excluded. All n values represent individual mice. All data were shown as mean ± SEM. Data were considered to be significant if p < 0.05.

### Differential expression and gene overlap of snRNA-seq dataset

Differential expression analysis after dataset integration was performed on the “RNA” assay after normalization, which is suggested in Satija lab’s website (https://satijalab.org/seurat/faq.html), using the Wilcoxon-rank-sum test of the Seurat package. The adjusted p-value was calculated using Bonferroni correction. Genes with |log(FC)| > 0.25 and adjusted p-value < 0.01 were selected as differentially expressed. To estimate the significance of overlap of two gene lists, overlapping p-value was calculated using the Fisher’s exact test of the GeneOverlap package (version 1.23.0) (https://rdrr.io/bioc/GeneOverlap/). The list of all genes expressed in a given cell type was used as the background. The lists of DAM- and DAA-upregulated genes were taken from supplementary table 2 in Keren-Shaul et al. ^48^ and from supplementary table 2 in Habib et al. ^49^, respectively.

### Enrichment analysis

Enrichment analysis was performed using Metascape ^87^ with default setting, which includes ontology catalogs of KEGG Pathway, GO Biological Processes, Reactome Gene Sets, and CORUM. Terms with –log10(FDR) > 2 were selected as significantly enriched. The heatmap of –log10(FDR) of all enriched terms was generated using Morpheus (https://software.broadinstitute.org/morpheus). Hierarchal clustering was performed with 1-Pearson correction and average agglomeration method.

